# Continuous muscle, glial, epithelial, neuronal, and hemocyte cell lines for Drosophila research

**DOI:** 10.1101/2023.01.18.524445

**Authors:** Nikki Coleman-Gosser, Shiva Raghuvanshi, Shane Stitzinger, Yanhui Hu, Weihang Chen, Arthur Luhur, Daniel Mariyappa, Molly Josifov, Andrew Zelhof, Stephanie E. Mohr, Norbert Perrimon, Amanda Simcox

**Author notes:** Equal contribution first author (Nikki Coleman-Gosser, Shiva Raghuvanshi, Shane Stitzinger).

## Abstract

Expression of activated Ras, Ras^V12^, provides Drosophila cultured cells with a proliferation and survival advantage that simplifies the generation of continuous cell lines. Here we used lineage restricted Ras^V12^ expression to generate continuous cell lines of muscle, glial, and epithelial cell type. Additionally, cell lines with neuronal and hemocyte characteristics were isolated by cloning from cell cultures established with broad Ras^V12^ expression. Differentiation with the hormone ecdysone caused maturation of cells from mesoderm lines into active muscle tissue and enhanced dendritic features in neuronal-like lines. Transcriptome analysis showed expression of key cell-type specific genes and the expected alignment with single cell sequencing data in several cases. Overall, the technique has produced in vitro cell models with characteristics of glia, epithelium, muscle, nerve, and hemocyte. The cells and associated data are available from the Drosophila Genomic Resource Center.

## INTRODUCTION

The use of cell cultures has been important for studying biological processes that are not easily accessible in whole organisms (Klein et al., 2022). A number of advances in mammalian cell cultures, for instance, development of 3D/organoid cultures (Rossi et al., 2018), improved genome editing tools to manipulate induced pluripotent stem cells (Hockemeyer & Jaenisch, 2016), and better optimized media formulations for recombinant protein expression (Ritacco et al., 2018) have further enhanced the utility of mammalian cell culture systems. These advances are accompanied by the availability of several distinct mammalian cell lines derived from different tissue types. Similarly, the use of insect cell lines also complements whole organismal studies and helped to illuminate many aspects of insect cell biology (Luhur et al., 2019) including development (Sato & Siomi, 2020), immunity (Chen et al., 2021; Goodman et al., 2021), host-pathogen relationships (Smagghe et al., 2009), in addition to biotechnological applications (Hong et al., 2022).

Fruit fly (*Drosophila melanogaster*) cell cultures are among the most widely used invertebrate cell cultures (Luhur et al., 2019). *Drosophila* cell lines are relatively homogenous, and highly scalable for both biochemical and high-throughput functional genomic analyses (Baum & Cherbas, 2008; Debec et al., 2016; Mohr, 2014; Viswanatha et al., 2019; Zirin et al., 2022). These features underlie their status as an important workhorse for scientific discovery in organismal development and as models for human disease. There are approximately 250 distinct *Drosophila* cell lines housed by the *Drosophila* Genomics Resource Center (DGRC)(Luhur et al., 2019). The majority of these cell lines, initially established by independent laboratories worldwide, were donated to the DGRC. A subset of 25 of these lines was subjected to transcriptome analysis, with the results demonstrating that approximately half of the transcripts expressed by each of these lines were unique such that even cell lines derived from the same tissue had distinct transcriptomic profiles (Cherbas et al., 2011). Furthermore, the transcriptional profiles of several imaginal disc lines analyzed were found to match profiles of cells from distinct spatial locations in the respective discs (Cherbas et al., 2011). All lines exhibited transcript profiles indicative of cell growth and cell division, and not cellular differentiation, as expected for proliferating cells (Cherbas et al., 2011). Thus, the transcriptional profiles of several Drosophila cell lines provided a platform for subsequent analyses. For instance, a few examples of the impact of this work include research into better understanding crosstalk between signaling pathways (Ammeux et al., 2016), exploring transcription factor networks (Rhee et al., 2014), establishing small RNA diversity (Wen et al., 2014), characterizing signaling pathways (Neal et al., 2019), nucleosomal organization (Martin et al., 2017) among multiple other utilities reviewed extensively (Cherbas & Gong, 2014; Luhur et al., 2019).

Over two-thirds of the *D. melanogaster* cell lines listed in the DGRC catalog were derived from whole embryos and the remainder are from various larval imaginal discs, the larval central nervous system, larval hemocytes, or adult ovaries. The potential of cells from these different sources to differentiate into adult cell types is not known. However, temporal transcriptional profiling of the Ecdysone response of 41 cell lines (Stoiber et al., 2016) provided evidence that cell lines exhibited varying levels of ecdysone sensitivity and potential for cellular differentiation, suggesting the possibility of developing cell-type specific cell lines with the capacity to differentiate.

As well as having unknown cellular origins, most Drosophila cell lines arose spontaneously, and the time needed to develop a continuous cell line was often protracted. In contrast, expression of activated Ras, Ras^V12^, using the Gal4-UAS system, resulted in the rapid and reproducible generation of continuous cell lines from primary embryonic cultures (A. Simcox et al., 2008). The Ras-method was used to develop an array of mutant cell lines by using appropriate genotypes to establish the primary cultures (Kahn et al., 2014; Lee et al., 2015; Lim et al., 2016; Nakato et al., 2019; A. A. Simcox et al., 2008). To date all lines have been generated using ubiquitous expression of *UAS-Ras* with *Act5C-Gal4* and therefore the cell type in a given line is unknown.

Here we describe a second-generation version of the Ras-method in which Ras^V12^ expression is restricted to a lineage by using tissue-specific Gal4 drivers. This genetic ‘dissection’ provides only the targeted cells with the survival and proliferation advantage conferred by Ras^V12^ expression (A. Simcox et al., 2008). As we show, the approach has been successful and resulted in the generation of cell lines with glial, epithelial and muscle characteristics. Lines generated by broad Ras^V12^ expression should also include those of specific cell types and by using single cell cloning and cell type characterization (marker gene expression and RNAseq) we identified lines with neuronal and hemocyte characteristics. Collectively these cell lines provide in vitro models for five different cell types and are expected to be a valuable resource for high throughput and biochemical approaches, which rely on large numbers of homogeneous cells.

## RESULTS

Primary cultures were established from embryos in which *UAS-Ras*^*V12*^ expression was restricted to glial, tracheal epithelial, and mesodermal cells using lineage-specific Gal4 drivers (Tables 1, S1). A subset of continuous cell lines derived from each type of primary culture was analyzed with regards to cell morphology, the presence of proteins characteristic of specific cell types, and other attributes (Table 1). We also analyzed lines with neuronal- or hemocyte-like characteristics that were cloned from parental lines resulting from ubiquitous expression of *UAS-Ras*^*V12*^ (Tables 1, S1). We further analyzed the cell lines by RNAseq. As expected, the transcriptomes of the new cell lines are distinct from those of existing cell lines (Cherbas et al., 2011) (Fig. S1) and new cell lines derived from the same Gal4 driver cluster with one another (Fig. S2). Moreover, comparison of differentially expressed (DE) genes with RNAseq data from single-cell RNAseq data (Li et al., 2022) (Table 2) or with known cell type-associated transcription factors (Fig. S3) reveals that these cells express genes characteristic of specific cell types. The results of our detailed characterization are described according to cell type in the sections below.

**Table 1.** Cell lines analyzed.

| Tissue-type alignment | Genotype | Lines analyzed* |
| --- | --- | --- |
| Glial | *repo-Gal4; Ras<sup>V12</sup>; brat<sup>dsRNA</sup>* | Rbr6 (parental)<br>Rbr6-2<br>Rbr6-4<br>Rbr6-F9 |
| Epithelial | *btl-Gal4; UAS-p35; UAS-Ras<sup>V12</sup>*<br>*btl-Gal4; UAS-p35; attP; UAS-Ras<sup>V12</sup>* | Btl3 (parental)<br>Btl7 (parental)<br>Btl8 (parental) |
| Muscle | *24B-Gal4; attP; UAS-Ras<sup>V12</sup>*<br>*24B-Gal4; UAS-GFP; attP; UAS-Ras<sup>V12</sup>* | 24B5 (parental)<br>24B5-B8<br>24B5-D8<br>24BG1 (parental)<br>24BG1-F3**<br>24BG1-G1** |
| Neuronal | *Act5C-GeneSwitch-Gal4; UAS-GFP; attP; UAS-Ras<sup>V12</sup>* | ActGSB-6<br>ActGSI-2 |
| Blood | *Act5C-GeneSwitch-Gal4; UAS-GFP; attP; UAS-Ras<sup>V12</sup>* | ActGSI-3 |

\* Clones unless indicated. \*\* Do not differentiate into active muscle.

**Table 2.** Identification of cell type by comparison of the top differentially expressed genes in cell lines with top marker genes from Fly Cell Atlas (FCA) scRNAseq datasets.

| Cell line name | Cell type from scRNAseq | log2 fold enrichment | Enrichment P value | scRNAseq dataset |
| --- | --- | --- | --- | --- |
| ActGSI-3 | hemocyte | 5.122 | 1.00E-25 | Whole body |
| ActGSI-2 | leg muscle motor neuron | 2.479 | 5.79E-03 | Whole body |
| ActGSB-6 | adult ventral nervous system | 2.8 | 7.56E-04 | Whole body |
| Rbr6-F9 | adult glial cell | 3.063 | 8.13E-05 | Whole body |
| Rbr6-4 | cell body glial cell | 2.8 | 7.56E-04 | Whole body |
| Rbr6-2 | adult reticular neuropil associated glial cell | 3.063 | 8.13E-05 | Whole body |
| Btl3 | adult tracheal cell | 2.063 | 2.61E-06 | Whole body |
| Btl8 | adult tracheal cell | 1.326 | 9.72E-03 | Trachea |
| Btl7 | adult tracheal cell | 1.849 | 8.81E-04 | Oenocyte |
| 24B5-B8 | muscle cell | 2.537 | 2.93E-06 | Male reprod. glands |

Analysis using the Drosophila RNAi Screening Center's single-cell Database (DRscDB), all datasets are 10x genomics.

### Glial-lineage cell lines

Repo is expressed exclusively in glial cells (Xiong et al., 1994). A *repo-Gal4* driver that recapitulates Repo expression was used to express *UAS-Ras*^*V12*^ (Ogienko et al., 2020; Sepp et al., 2001). This led to robust production of primary cultures however these failed to survive beyond early passages (Table S1). To counter potential cell death or modulate growth signaling, additional genotypes were tested including co-expression of *UAS-transgenes* encoding the P35 baculovirus cell survival factor, dsRNAs targeting tumor suppressors, or the Gal4 inhibitor Gal80^ts^ (Table S1). Co-expression of a *UAS-brat*^*dsRNA*^ or expression of *tub-Gal80*^*ts*^ each produced a single line of cells that could be propagated for extended passages however the latter line was difficult to maintain and eventually lost (Table S1). The *repo-Gal4: UAS-brat*^*dsRNA*^; *UAS-Ras*^*V12*^ (Rbr6) line has been passaged more than fifty times. The parental Rbr6 line and three clonal derivatives (Rbr6-2, Rbr6-4 and Rbr6-F9) have an elongated morphology and stained positive for Repo (Table 1; Figs 1, S4, S5A). A few cells expressed neuronal markers (Fig. S5A; Table S2). To induce differentiation, we gave cells two 24 h ecdysone treatments separated by 24 h to approximate the pulses of ecdysone during the larval to pupal transition. Cells from each of the clones survived treatment with ecdysone suggesting they are of adult type, two clones showed morphological changes and formed a network, and all continued to express Repo (Fig. 1, S6).

**Fig. 1.**
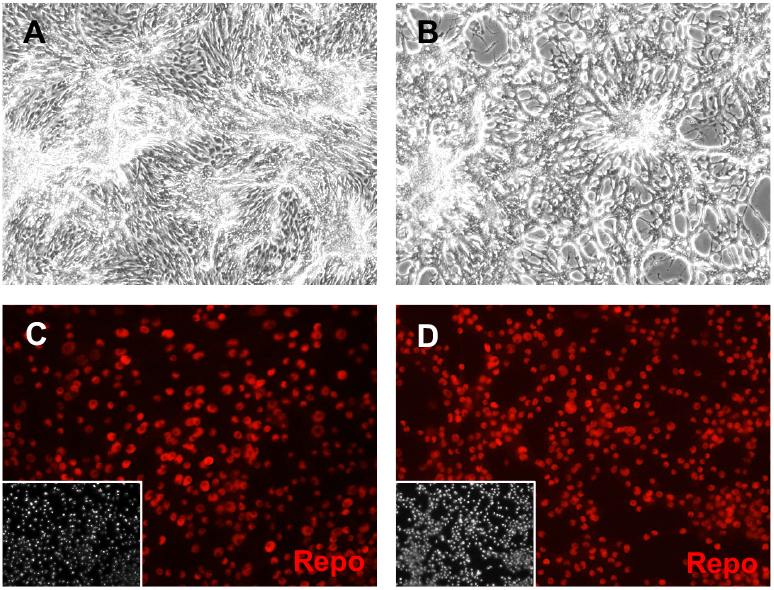
Glial Clone Rbr6-2 cells express Repo. Cells were grown in plain medium (A,C) or treated with ecdysone (B,D). (A,B) After ecdysone treatment, cells make a lace-like network. (C,D) Cells express Repo with or without ecdysone treatment. Inset: DAPI, DNA

The results of RNAseq analysis revealed that the three Rbr6 clones have very similar expression patterns (Fig. S2). In addition, their DE gene signatures are also a close match to gene signatures of glial cells as identified by single-cell RNAseq (Table 2) and to glial-associated genes reported in the literature. For example, *zydeco (zyd)*, which encodes a potassium-dependent sodium/calcium exchanger, is upregulated in all three clones, consistent with the literature (Featherstone, 2011; Zwarts et al., 2015), and *gcm2*, a transcription factor, is upregulated in two clones (Fig. S3). These data suggest the Rbr6 clones will be a useful in vitro source of glial cells.

### Tracheal epithelium-lineage cell lines

Breathless is expressed in the tracheal epithelium and a *btl-Gal4* driver was used to express *UAS-Ras*^*V12*^ (Shiga et al., 1996). Patches of cells with epithelial morphology proliferated in primary cultures and several continuous lines were generated (Tables 1, S1). We were unable to derive clones of these using dilution or selection methods, which were successful for other cell types. Correspondingly, three parental lines were examined; Btl3, Btl7, and Btl8 (Table 1). All showed expression of the epithelial marker Shotgun/E-Cadherin (Shg/Ecad) and two grew in a squamous epithelial sheet with Ecad expression at the cell periphery (Figs 2, S5B, S7). Treatment of the squamous epithelial cells (Btl3, Btl7) with ecdysone caused aggregation and formation of large multicellular clusters (Figs 2, S8).

**Fig. 2.**
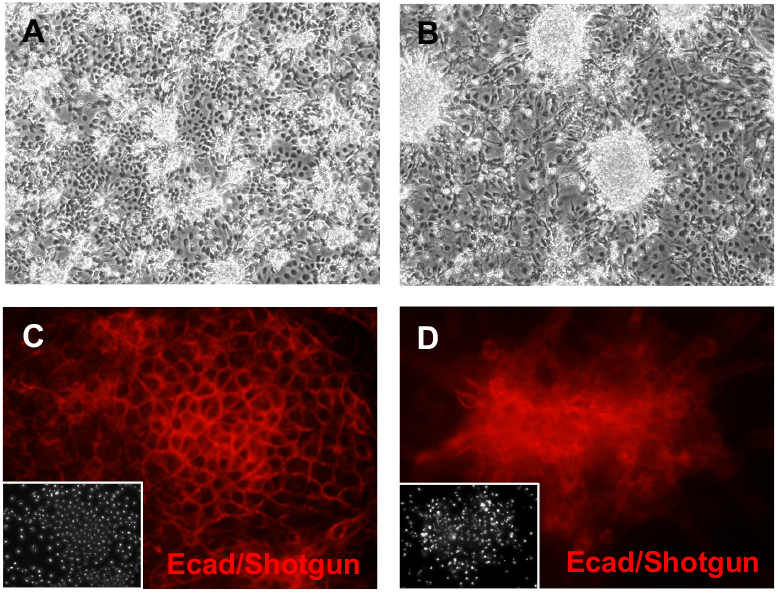
Tracheal lineage cells of line Btl3 express the epithelial cadherin Ecad/Shotgun. Cells were grown in plain medium (A,C) or treated with ecdysone (B,D). (A) Cells form a squamous epithelial sheet. (B) Ecdysone treated cells form clusters that pile up. (C) Control cells form a cell sheet with small cell clusters and expressed Ecad at the cell boundaries. (D) Ecdysone treated cells form large multicellular clusters that expressed Ecad.

RNAseq data analysis comparing the top up-regulated genes in the Btl cell lines with scRNAseq datasets revealed that two of the three parental lines (Btl3 and Btl8) closely match the signatures of the adult tracheal, a network of epithelial tubules (Table 2) and Btl3 expresses *trachealess (trh)* a master regulator of tracheal identity (Wilk et al., 1996)(Fig. S3). Overall, the morphological and molecular characteristics of the lines are consistent with an epithelial cell type of tracheal origin.

### Mesodermal-lineage cell lines

The *24B-Gal4* driver is an insertion in *held out wings (how)* and is expressed in mesoderm and muscle cells (Brand & Perrimon, 1993; Zaffran et al., 1997). Expression of *UAS-Ras*^*V12*^ with *24B-Gal4* readily produced continuous lines (Tables 1, S1). Four clones (24B5-B8, 24B5-D8, 24BG1-F3, and 24BG1-G1) derived from two parental lines (24B5 and 25BG1) were analyzed in more detail (Table 1). The cells had a bipolar shape and expressed mesoderm markers including Twist and Mef2 (Figs 3, S5C, S9). When treated with ecdysone, cells from both parental lines and clones 24B5-B8 and 24B5-D8 elongated, formed a network, and expressed Myosin heavy chain (Mhc) (Figs 3, S10). There was also extensive cell lysis. Beginning two days after the second ecdysone treatment, the cells began to contract spontaneously.

**Fig. 3.**
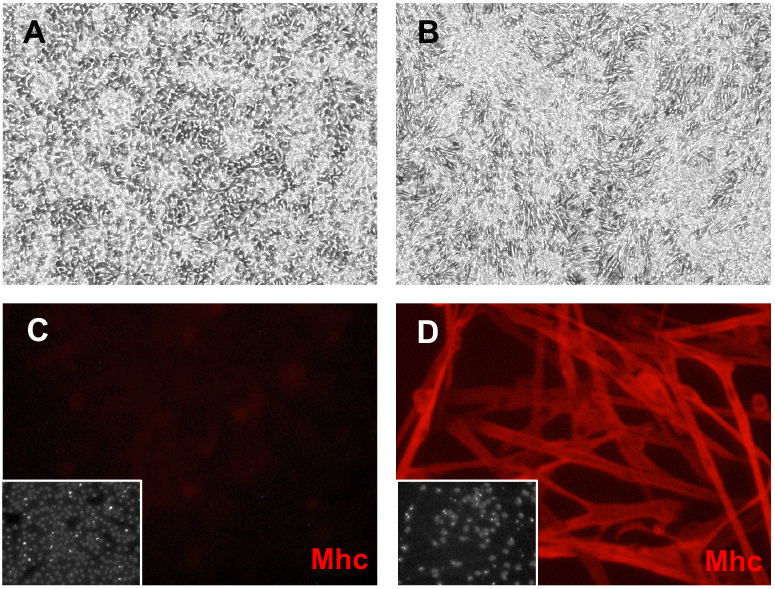
Mesodermal lineage cells of Clone 24B5-B8 express. Myosin heavy chain after differentiation. Cells were grown in plain medium (A,C) or treated with ecdysone (B,D). (A) Cells have a bipolar shape. (B) Ecdysone treated cells elongate and contract. (C) Control cells do not express Mhc. (D) Ecdysone treated cells express the muscle marker Mhc. Inset: DAPI, DNA.

Contraction of cells from the 24B5 parental line and the two derivative clones (24B5-B8 and 24B5-D8) was visible in real time (Movies 1, 2), whereas contraction of parental line 24BG1 cells was much slower and visualized more clearly in time-lapse (Movies 3, 4). The clones 24BG1-F3, and 24BG1-G1 underwent morphological change but did not express Mhc or contract (Figs S10, S11). In later passages, the 24BG1 parental line also lost expression of Mhc and the ability to contract (S10). This highlights the importance of using early passage cells and avoiding extended passaging that could alter the phenotypic (and genotypic) characteristics of the cells.

The RNAseq analysis for these cell lines, which as for the others was performed on undifferentiated cells, does not result in as clear a similarity to an expected cell type as was observed for Rbr6 or Btl cells. However, we did find that undifferentiated 24B5-B8 cells express high levels of the transcription factors *nautilus (nau)* and *twist (twi)* (Fig. S3; Table 2), and high levels of *myoblast city (mbo)*, which encodes an unconventional bipartite GEF with a role in myoblast fusion (Erickson et al., 1997). The capacity of these mesoderm-derived cell lines to differentiate into active muscle shows that the cells are muscle precursors and thus should be a useful reagent to analyze muscle physiology and development.

### Neuronal-like cell lines

We expressed *UAS-Ras*^*V12*^ with the pan-neural drivers *scratch-Gal4* and *elav-Gal4*, however none of the primary cultures resulted in continuous cell lines (Table S1). In previous work, we made primary cultures from embryos with ubiquitous expression of *UAS-Ras*^*V12*^ using the *Act5C-Gal4* driver (A. Simcox et al., 2008). The cells growing in these cultures included neuronal cells (A. Simcox et al., 2008). Here we used an *Act5C-GeneSwitch-Gal4* driver to express *UAS-Ras*^*V12*^. GeneSwitch-Gal4 is only active in the presence of the drug, RU486/mifepristone, which provides the advantage of being able to regulate Ras^V12^ expression (Nicholson et al., 2008; Osterwalder et al., 2001). Several continuous lines were generated (Table S1). Clones derived from two of these (ActGSB-6 and ActGSI-2)(Table 1) were positive for the neuronal marker, HRP (Figs 4, S5D, S12, S13). After differentiation with ecdysone, expression of Futsch/MAPB1 (Hummel et al., 2000) and Fas2 (Mao & Freeman, 2009) was enhanced and revealed axonal-like outgrowths from the cells (Figs 4, S13). Differentiated cells also showed enhanced expression of Elav, which is commonly used as a marker for postmitotic neurons (Figs 4, S10) (Robinow & White, 1991). Elav is also expressed transiently in glial cells and proliferating neuroblasts (Berger et al., 2007) however the cells were negative for the glial marker Repo (Table S2).

**Fig. 4.**
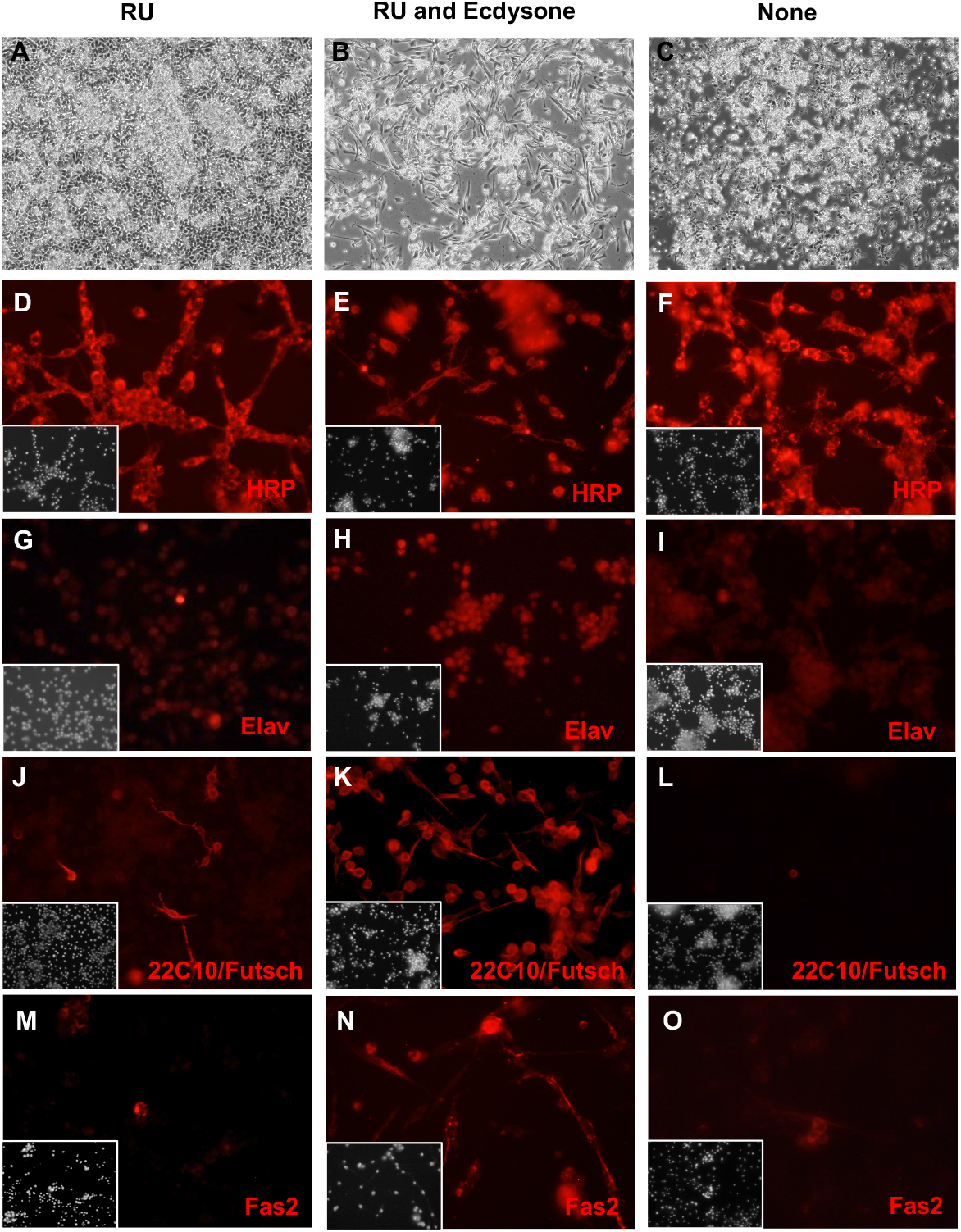
Neuronal-like clone ActGS-I2 expresses neuronal markers. ActGSI-2 cells were grown in three conditions: RU486 (A,D,G,J,M); RU486 and ecdysone (B,E,H,K,N), or with no additives (C,F,I,L,O). RU486/mifepristone is required for GeneSwtch-Gal4 activation, transgene expression, and cell proliferation. (A) In the growing condition, cells reach confluence and continue to grow by piling up. (B) After ecdysone treatment cells undergo morphological change including extending dendritic-like processes. (C) In the quiescent state (no RU), cells do not proliferate and fail to reach confluence. (D,E,F) Cells in all conditions are positive for HRP. (G,H,I) Expression of Elav, is elevated after ecdysone treatment (H). (J,K,L) Expression of Futsch/MAP1B-like protein (recognized by antibody 22C10) is elevated after ecdysone treatment (K). (M,N,O) Fas2 neural-adhesion protein. Cells show elevated expression after ecdysone treatment (N).

RNAseq analysis revealed that many neuronal genes are up-regulated in these cell lines, including *Glutamic acid decarboxylase 1* (*Gad1*), *slowpoke* (*slo*), *5-hydroxytryptamine (serotonin) receptor 1A* (*5-HT1A*), *Protein C kinase 53E* (*Pkc53E), Diuretic hormone 31 Receptor* (*Dh31-R*), and *straightjacket (stj)*. In addition, comparison of the top up-regulated genes in these cells to marker genes from scRNAseq data identifies a cell type of neuronal origin as the best match (Table 2). The cells should be a useful source of neuronal cells.

### Hemocyte-like cell line

Cells of clone ActGSI-3 derived from the ActGSI parental line *(UAS-Ras*^*V12*^ expression with *Act5C-GeneSwitch-Gal4*; Tables 1, S1) have a distinct morphology and an unusual growth cycle in which cells initially divide in floating clusters and then disaggregate as single cells on the surface of the tissue-culture dish (Fig. 5). Interestingly, the cells express the hemocyte marker Hemese (Fig. 5)(Kurucz et al., 2003). They are also positive for HRP, but not other neuronal markers (Fig S5E).

**Fig 5.**
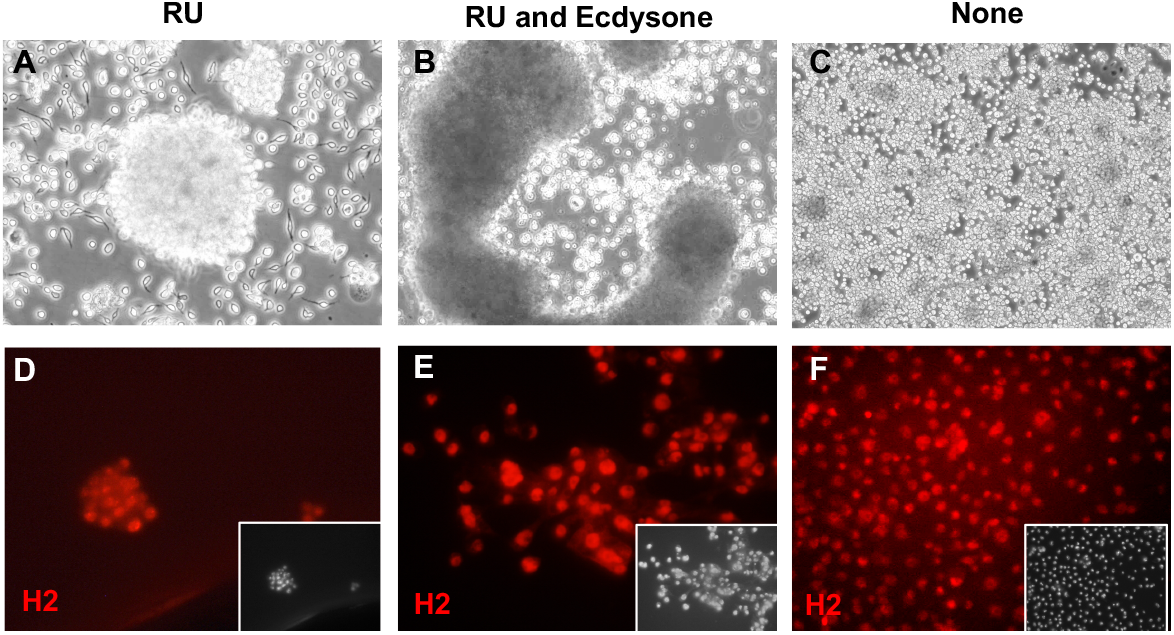
Hemocyte-like Clone ActGS-I3 morphology and marker expression. Cells were grown in three conditions: RU486 (A,D); RU486 and ecdysone (B,E), or with no additives (C,F). (A) In the growing condition, cells are seen as individual round/bipolar cells or as floating clusters of multiple cells. (B) After ecdysone treatment cells formed large aggregates and there was cell lysis. (C) In the quiescent state (no RU), individual round cells are seen. (D,E,F) Cells in all conditions express the hemocyte cell marker Hemese, as recognized by the antibody H2. Inset: DAPI, DNA.

RNAseq analysis demonstrated that many hemocyte genes are upregulated in these cells, including *serpent (srp), Hemese (He), eater, u-shaped (ush), Cecropin A2 (CecA2)*, and *Cecropin C* (CecC). Comparison of top up-regulated genes with scRNAseq data showed that the cells have a strong match to the top marker genes of hemocytes (Table 2).

### Growth, karyotype, and transfection efficiency of cell lines

We determined the doubling time of 13 cell lines and clones using growth curves (Fig. S11; Table 3). Most had doubling times within a range of approximately 20-40 h (Table 3). The hemocyte-like clone ActGSI-3 was an outlier with a much longer doubling time of 70 h (Table 3). We determined the karyotype of eight cell lines and clones. In keeping with previous findings for Ras^V12^ expressing cell lines, most were diploid, or near diploid (A. Simcox et al., 2008) (Fig. S15; Table 3). The ActGSI-3 clone was mixoploid with chromosome numbers ranging from 11 to 18 (Fig. S15).

**Table 3.**
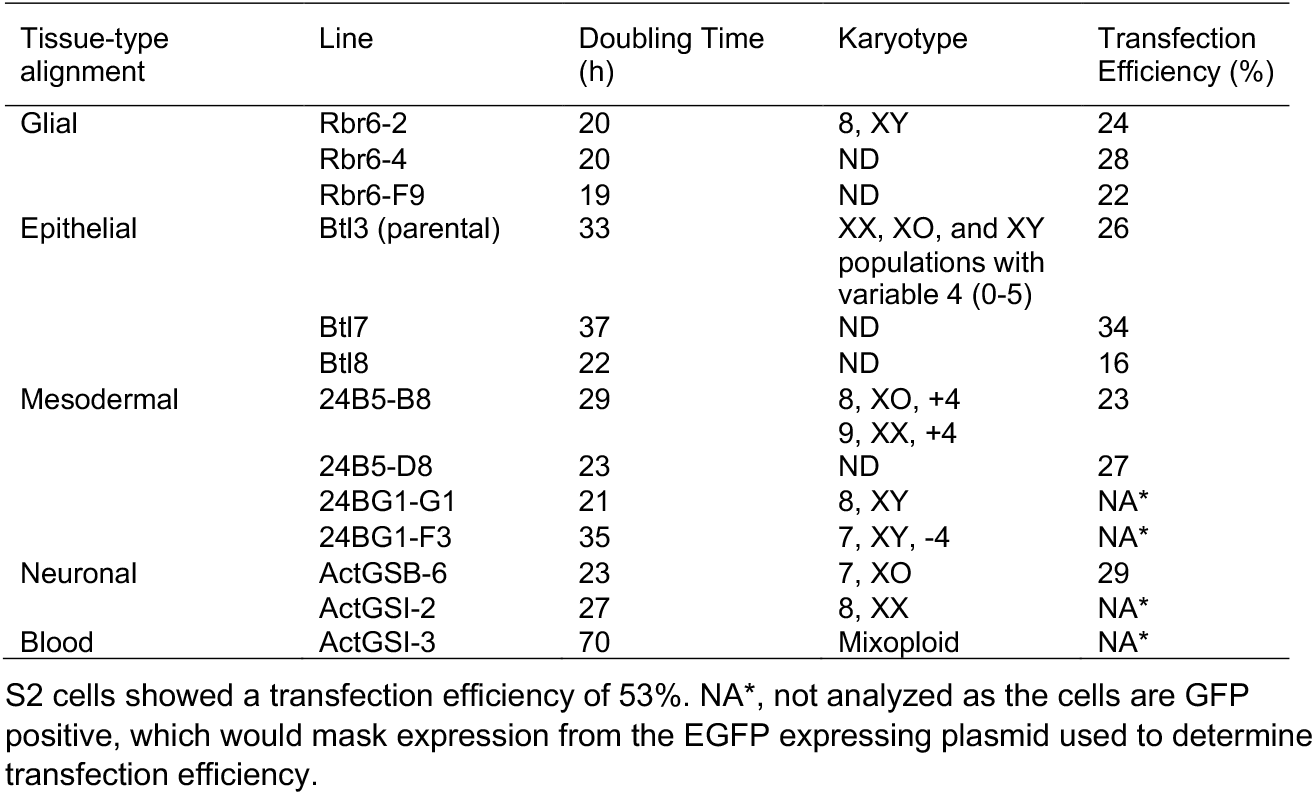
Doubling time, karyotype, and transfection efficiency of select cell lines.

Nine parental and clonal lines were transfected with an Act5C-EGFP plasmid and the fraction of GFP positive cells was determined after 48 hours. Cells from all lines tested could be transfected. The range of efficiency was from 16% to 34% with most lines showing transfection of approximately one quarter of the cells (Table 3). Similarly treated, cells from the S2 line showed an efficiency of 53%.

## DISCUSSION

Expressing activated Ras, Ras^V12^, in primary cells provides a growth and survival advantage and leads to the rapid and reliable generation of continuous cell lines—the so-called Ras method (A. Simcox et al., 2008). In a second-generation version of the Ras-method, we found that restricting Ras^V12^ expression with lineage-specific Gal4 drivers gave the targeted cells a competitive advantage and produced continuous lines with expected cell-type specific phenotypes. With this approach we produced glial, epithelial, and muscle cell lines using the *repo-, btl-*, and *24B/how-Gal4* drivers, respectively.

In theory, the approach could be used to produce cell lines corresponding to any cell type for which there is an appropriate Gal4 driver. However, for the muscle lineage, we also tried to derive lines with *Mef2-Gal4*, but no continuous lines were produced (Table S1). Mef2 regulates muscle differentiation and is expressed in muscle progenitors and differentiated muscle suggesting *Mef2-Gal4* would be a good candidate for deriving cell lines (Bour et al., 1995; Gossett et al., 1989; Lilly et al., 1995; Ranganayakulu et al., 1995). Similarly, we failed to produce lines from the pan-neuronal drivers *elav-Gal4* and *scratch-Gal4* (Table S1). The failure to derive continuous cell lines from these drivers suggests that it may be necessary to test multiple Gal4 lines for a given lineage.

*repo-Gal4* is a pan-glial driver and many primary cultures expressing Ras^V12^ with this driver reached confluence and could be passaged 1-6 times. We tested different genotypes to determine if the success rate could be improved by modulation of Ras^V12^ expression (co-expression of the Gal4 inhibitor Gal80ts), co-expression of the p35 baculovirus survival factor, or growth stimulation by downregulation of tumor suppressors (dsRNA for *warts* or *brat)*. One line, also harboring a Gal80ts transgene, reached passage 25; however, the line was unstable and in early passages the cells variably lost Repo expression and changed morphologically. The one continuous glial line generated expresses a transgene that targets the tumor suppressor, *brat, (repo-Gal4; UAS-Ras*^*V12*^; *UAS-brat*^*dsRNA*^). Given a single success, it is not clear if down regulation of *brat* contributed to derivation of the line. Moreover, there is no evidence that these genotypic variations enhanced cell line generation with other drivers, as primary cultures expressing Ras^V12^ without modulation or a survival factor produced lines with similar success rates for the *btl-Gal4* or *24B/how-Gal4* drivers (Table S1).

As with all types of tissue culture, best practices involve maintaining frozen aliquots of cell lines at relatively low passage numbers. Aliquots of cells from the lines and clones described here, on which RNAseq was performed, have been archived at similar passage numbers as those used for the RNAseq analysis. This will allow users to start experimentation with the lines in a known state. The importance of this is exemplified by line 24BG1, which lost the ability to contract and express the muscle protein Mhc over the course of 15 passages (Fig. S10).

The mesodermal, neuronal, and glial cells represent in vitro counterparts of the tissues of origin that can be used for studying development and physiology in an accessible and reproducible system. The mesodermal cells that differentiate into active muscle will allow investigation of muscle fusion, as the cells are multinucleate, as well as muscle physiology and function. For example, the cells contract spontaneously and in apparent waves (Movies 1-3); however, the mechanism for stimulation (if any) and regulation have not been investigated and may cast light on in vivo processes. Given a variety of cell types, it will also be interesting to examine cell form and function in co-cultures, for example, of glia and neurons.

The method and the cells will be useful for generating disease models. New lineage specific lines could be generated in the desired mutant background by establishing primary cultures from embryos in which only the mutant genotype expresses Ras^V12^ giving these cells a growth and survival advantage (A. A. Simcox et al., 2008). Derivative lines should include those of the desired cell type and genotype. Alternatively, the existing cell lines could be edited using CRISPR, or insertion of transgenes using the attP site that most lines and clones contain (Table S1)(Bateman et al., 2006; Manivannan et al., 2015).

The cells with epithelial morphology derived from the tracheal lineage (Btl3 and Btl7) will provide good models for investigating assay conditions that promote polarization and 3D cell interactions that could allow the cells to manifest a more complex tissue architecture. In keeping with this possibility, treating these cells with ecdysone to induce differentiation showed cell clumping suggestive of a multicellular structure (Fig. S8).

RNAseq analysis of cells from the I3 cell line showed a striking similarity to hemocytes and the cells may be a good model for studying immunity (Table 2). The cells lyse after ecdysone treatment suggesting they are of embryonic origin (Fig. 5). The cells have an unusual growth cycle transitioning between clusters and single cells (Fig. 5). This may recapitulate the sessile (subepidermal clusters) and circulating hemocytes of the larva (Leitao & Sucena, 2015).

The most significantly up-regulated marker genes in each cell line are significantly enriched for top marker genes from expected cell types based on the single-cell RNAseq data from Fly Cell Atlas in most cases. This indicates the potential value of these cell lines as corresponding in vitro models for studying these cell types. While the cells will prove to be valuable models, it should be noted that even those showing a clear differentiated phenotype exhibit unexpected patterns of gene expression. For example, some cells in the mesodermal clone, 24B5-B8, are positive for HRP (Fig. S3) and the two neuronal-like lines express a mesodermal marker, Twist (Fig. S3). This anomalous gene expression is likely to be an effect of Ras activation on downstream pathways and genes. Ras/MAPK has a key role in muscle cell determination (Buff et al., 1998; Carmena et al., 2002; Halfon et al., 2000) and activates downstream muscle determination genes.

The cells will have value for both low- and high-throughput approaches, including genetic or compound screens for which screening in the relevant cell type will result in identifying targets that are more likely to be of physiological relevance. Most of the cells have an attP-flanked cassette (Table 1), which makes them amenable to insertion of transgenes such as reporters by Recombination Mediated Cassette Exchange (RMCE) (Bateman et al., 2006; Manivannan et al., 2015). Moreover, cells competent for RMCE can be modified by stable expression of Cas9 and then used for genome-wide CRISPR pooled screening. With this approach, a library of single guide RNAs (sgRNAs) are integrated at RMCE sites (Viswanatha et al., 2019; Viswanatha et al., 2018). This generates a pool of cells, each with a different sgRNA, that can be subjected to a screen assay. Results are identified by PCR amplification of inserted sgRNAs followed by next-generation sequencing (NGS) to detect sgRNAs that are enriched or depleted in the experimental cell pool as compared with a control. To date, pooled CRISPR screens in Drosophila have only been performed in S2 cells, which have hemocyte-like features. The availability of new cell lines with muscle, glial, and epithelial characteristics will enable screens designed to interrogate biological processes specific to these cell types.

There are hundreds of Drosophila cell lines; however, the number corresponding to known cell types is low. This is due in part to the lack of a method for generating cell lines from specific tissues. We expect that the method described here, using restricted expression of Ras^V12^, will be a tractable approach for investigators to generate lines of cell types of interest. Single cell cloning followed by cell characterization (immunohistochemistry and RNAseq) also proved to be a useful method to identify cell-type specific lines and this approach could identify additional valuable lines in the existing collection at the DGRC. In summary, we show that lineage-restricted Ras expression and cell cloning has produced a set of new cell lines that will be of immediate value for analyses in the five cell types they represent.

## MATERIALS and METHODS

### Fly Stocks

The following fly stocks were used to create primary cell lines: Gal4 drivers: 24B/how-Gal4, w[*]; P(w[+mW.hs]=GawB)how[24B] (BL 1767); repo-Gal4, P(GAL4)repo (BL 7415); btl-Gal4, P(GAL4-btl.S)3-2 (BL 78328); Act5C-GeneSwitch-Gal4, P(UAS-GFP.S65T)Myo31DF[T2]; P(Act5C(-FRT)GAL4.Switch.PR)3 (BL 9431). Transgenes: UAS-Ras^V12^ (3), P(w[+mC]=UAS-Ras85D.V12)TL1 (BL 64195); UAS-Ras^V12^ (2), P(w[+mC]=UAS-Ras85D.V12)2 (BL 64196); UAS-Ras^V12^ with RMCE site (3), P(w[+mC]=UAS-Ras85D.V12)TL1, P(w[+mC]=attP.w[+].attP)JB89B (BL 64197); UAS-GFP nuclear, P(UAS-GFP.nls)14 (BL 4775); brat^dsRNA^, P(y[+t7.7] v[+t1.8]=TRiP.HMS01121)attP2 (BL 34646); UAS-p35 baculovirus death inhibitor, P(w[+mC]=UAS-p35.H)BH1 (BL 5072) and Gal80ts, w[*]; P(w[+mC]=tubP-GAL80[ts])20 (BL 7019).

### Primary Cultures and Passaging

Primary cultures were established using the detailed method described in (Simcox, 2013) except that two embryo collections were made each day; one from approximately 9AM to 5PM at room temperature and one overnight at 17°C. The shorter collection tended to produce better primary cultures likely because on average the embryos were younger. Primary cultures were passaged once they reached about 70% confluence using trypsin to dislodge the cells (Simcox, 2013). Early passages were set up at low dilution (1 in 2). Established lines differ in their growth characteristics (Table 3) however most are passaged every 5-7 days at a dilution of 1 in 4.

### Cell Cloning

For puromycin selection, 2-6×10^5^ cells in a 35 mM well were transfected with 0.4ug of DNA encoding a puro resistance plasmid (pCoPURO, Addgene #17533) using Effectene Transfection Reagent (Qiagen). After 24 h, cells were selected with puromycin at 0.5-2.5ug/mL for 5 days. After 2-4 weeks, colonies were isolated and expanded. For dilution cloning, cells were seeded into a 96-well plate at a concentration of 0.5-1 cell/well in 100uL conditioned media (Housden et al., 2015).

### Hormone treatment

To simulate the major pulse of ecdysone at the larval to pupal transition, cells were treated with two 24 h doses of *β*-ecdysone (Sigma 5289-74-7) at 1 ug/ml separated by 24 h in non-supplemented medium.

### Immunohistochemistry

Cells were fixed with 4% paraformaldehyde (Electron Microscopy Sciences) for 15 min or 3.5% formaldehyde (Sigma) for 30 min at room temperature, and then rinsed twice with 0.1% Tween-20 in PBS (PBS-T). Cells were permeabilized (0.2% Triton X-100 in PBS) for 10 min at room temperature. Cells were blocked (5% BSA in PBS-T) for 30 min at room temperature and incubated with diluted primary antibodies overnight at 4°C. Cells were washed three times with PBS-T and incubated with diluted secondary antibodies in blocking buffer for 1 h at room temperature or overnight at 4°C. Cells were washed three times with PBS-T and mounted in VectaShield with DAPI (Vector Laboratories). For the Dcad2 antibody, cells were fixed and processed as described in (Oda et al., 1994). The following primary antibodies and dilutions were used: HRP (rabbit polyclonal, Jackson ImmunoResearch 323-005-021, 1:500), 22C10 (mouse monoclonal anti-Futsch, Developmental Studies Hybridoma Bank, DSHB, 1:100), ELAV (rat monoclonal, DSHB 7E8A10), Repo (mouse monoclonal, DSHB 8D12, 1:100), FasII (mouse monoclonal, DSHB 1D4, 1:100), Twist (a gift from M. Levine, UC Berkeley, CA, guinea pig 1:500), MHC (mouse monoclonal, DSHB 3E8-3D3, 1:100), Dcad2 (rat monoclonal, DSHB, 1:100), and DMef2 (a gift from J. R. Jacobs (Vanderploeg et al., 2012), rabbit polyclonal, 1:500), H2 (mouse monoclonal, (Kurucz et al., 2003), 1:10). Cells were incubated with the following secondary antibodies at the indicated dilutions: Cy3-conjugated goat anti-mouse (Jackson ImmunoResearch 115-165-003, 1:1000), Cy3-conjugated goat anti-rat (Jackson ImmunoResearch 112-165-003, 1:1000), Cy3-conjugated goat anti-guinea pig (Jackson ImmunoResearch 106-165-003, 1:1000), Cy3-conjugated goat anti-rabbit (Jackson ImmunoResearch 111-165-045), and Alexa Fluor 488-conjugated donkey anti-rabbit (Invitrogen A-21206, 1:1000).

### Growth curve analysis

1-2×10^5^ cells were plated in a 12-well plate. Cells were counted from triplicate wells every 3 days over a 9-day period. Doubling time was calculated using log2 cell numbers (Roth, 2006).

### Karyotype analysis

Cells were grown to 50% confluence and incubated with 0.05ug /mL KaryoMAX (Gibco-ThermoFisher 15212012) for 3-18 h. The cells were trypsinized, rinsed with PBS, and resuspended in 5 mL of hypotonic solution (0.075M KCl/ 1.3mM sodium citrate) by dropwise addition. After a 1 h incubation, 4 drops of fresh cold fixative (3:1, methanol: acetic acid made fresh) were added, the tube was centrifuged, and the pellet was resuspended in 3 mL fixative and incubated for 10 min. Cells were centrifuged, resuspended in a small volume of fixative, and spotted onto slides by dropping from a height of 70cm to promote spreading. Slides were air dried and mounted in VectaShield with DAPI (Vector Laboratories).

### Transfection

Cells in a 6-well plate (approximately 70% confluent) were transfected with 0.4ug of an Actin5C-EGFP plasmid (pAc5.1B-EGFP, Addgene #21181) using Effectene Transfection Reagent (Qiagen). The fraction of GFP positive cells was scored after 48 h.

### RNA extraction and RNAseq

Cell cultures were grown and expanded in their respective media. All cell lines were cultured in Schneiders Drosophila Medium (Gibco Cat # 21720001), supplemented with 10% Fetal Bovine Serum (Cytiva Hyclone Cat SH30070.03). For Act5C-GS>Ras attP-GFP-LI-Clone 2, Act5C-GS>Ras attP-GFP-LI-Clone 3 and Act5C-GS>Ras attP-GFP-LB-Clone 6 cultures were grown in the same basal media supplemented with 10 nM of Mifepristone (Thermofisher Cat# H11001). Cultures were allowed to grow in T-25 flasks to become confluent before treatment with trypsin (Gibco Cat# 12604013) for 4 minutes to dislodge the cell monolayer from the growth surface. The cells were resuspended in 4 mL of their respective media and 1 mL of the cell suspension was collected for pelleting, followed by washing in 1X PBS, and then flash-freezing in liquid nitrogen. All cell samples were processed in triplicates.

Total RNA was isolated from the pellets using the TRIzol^™^ reagent (Life technologies (Ambion), Cat#:15596018) as per manufacturer’s instructions. The isolated total RNA was subjected to further purification using the RNeasy Mini Kit (Qiagen, Cat#74104) and the RNA post-cleanup was eluted in RNase-free water. The eluted total RNA was confirmed to have a A_260_/A_280_ ratio > 1.8 and RIN > 7.

Upon passing the quality control parameters, Illumina TruSeq libraries were constructed using TruSeq stranded mRNA HT kit (Illumina, Cat# RS-122-2103). Paired end sequencing was performed on an Illumina NextSeq 500 with a 150-cycle high output kits (Illumina, Cat# FC-404-2002).

### RNAseq Data Analysis

Raw data processing was performed using the STAR sequence aligner (https://github.com/alexdobin/STAR)(Dobin et al., 2013). Reads were aligned to the Drosophila genome and featureCounts were used to get gene counts from all samples into a count matrix for downstream analysis. A PCA plot was produced using heatmaply. FPKM values were calculated using fpkm(DEseq2) using gene length output by featureCounts. The reference genome used was FB2022_05, dmel_r6.48 (FlyBase)(Jenkins et al., 2022). Both raw sequencing reads and the count matrix were deposited in the NCBI Gene Expression Omnibus (GEO) database under the accession number GSE219105.

Each sample was compared against all other samples by using DESeq2 (https://bioconductor.statistik.tu-dortmund.de/packages/3.4/bioc/html/DESeq2.html) to determine differentially expressed genes (DE calling). The set of top DE genes for each cell line was compared with the top 100 markers in single-cell RNAseq datasets corresponding to cell types in the Fly Cell Atlas 10x datasets (Li et al., 2022). Enrichment analysis was conducted using the DRscDB tool to identify the Fly Cell Atlas cell type that matched closely to each cell line (Hu et al., 2021).

The RNAseq data for the cell lines described in this work were also compared with RNAseq datasets determined previously for 24 other Drosophila cell lines (Cherbas et al., 2011). The comparison was conducted by hierarchical clustering analysis using Pearson correlation co-efficient scores.

## Acknowledgements

We thank M. Levin, J. R. Jacobs, and D. Hultmark for antibodies and the Bloomington Stock Center for fly stocks. We thank Mikhail Kouzminov for help with data analysis.

## Competing Interests

The authors declare no competing or financial interests.

## Author Contributions

Conceptualization: A.S.; Methodology: A.S.; Formal Analysis: A.S., N.C, S.S., Y.H., W.C.; Investigation: A.S., N.C, S.R., S.S., M.J., A.L, D.M; Writing - original draft: A.S.; Writing - review & editing: N.C, S.S., S.E.M., N.P., Y.H., A.L., D.M., A.Z.; Supervision: A.S, S.E.M., N.P.; Funding acquisition: A.S., S.E.M., N.P., A.Z,.

## Funding

This work is supported by the National Institutes of Health (NIH Office of the Director R24 OD019847 to N.P., S.E.M. and A.S., and P40OD010949 to DGRC) the National Science Foundation (IOS 1419535 to A.S.), the Howard Hughes Medical Institute (N.P.), and a gift from Women & Philanthropy at the Ohio State University (to A.S.).

## Data availability

The raw and processed RNAseq datasets were deposited in the NCBI Gene Expression Omnibus (GEO) database under accession code GSE219105.

## Materials availability

All cell lines described here have been deposited to the Drosophila Genomics Resource Center (DGRC) at Indiana University. The lines will be available for distribution to the research community immediately after publication.

---

**Supplementary Figure 1:**
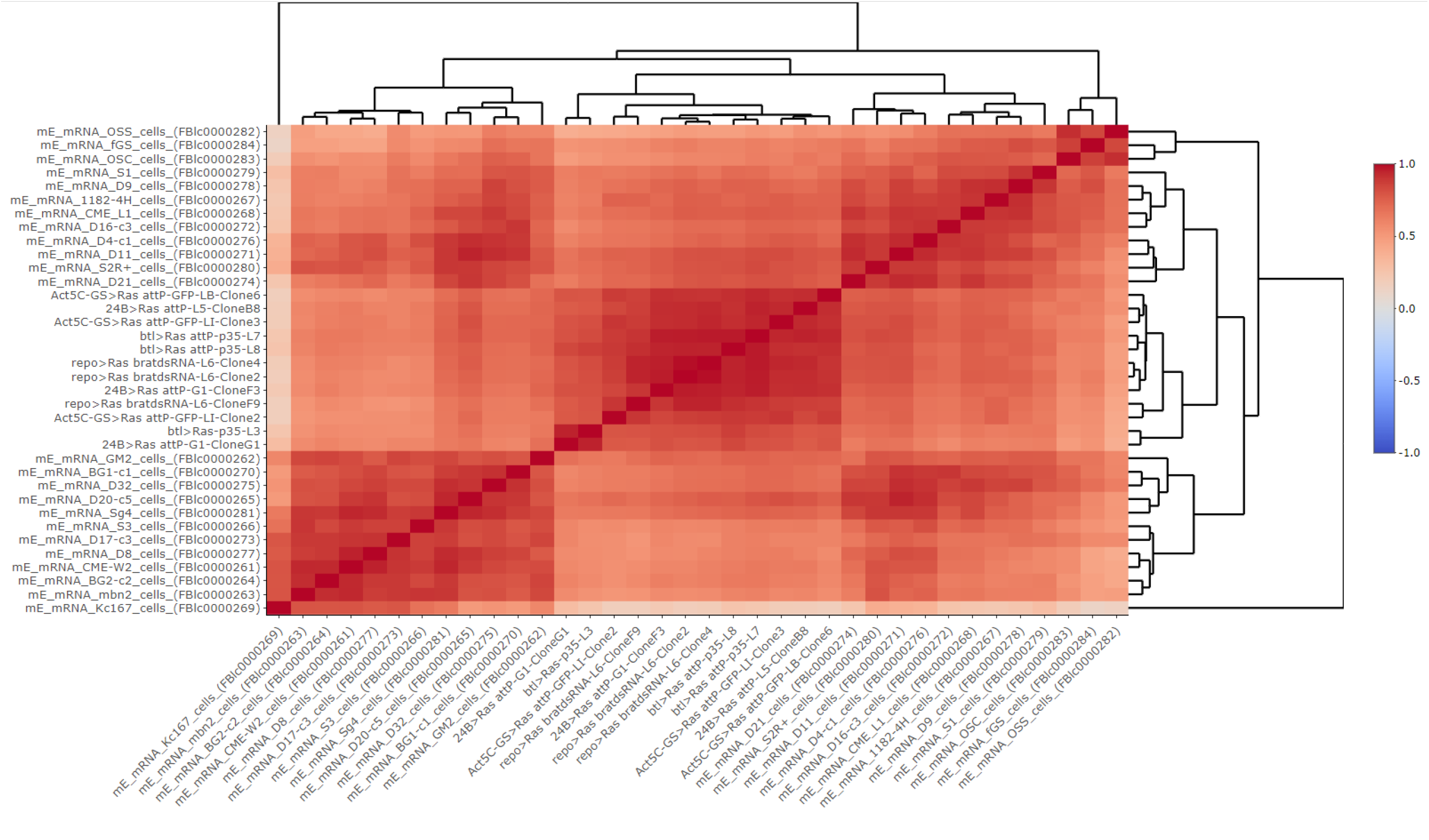
Comparison of lineage-restricted Ras cell lines with previously isolated Drosophila cell lines. RNAseq data for the cells described in this work were compared with RNAseq datasets determined previously for 24 other Drosophila cell lines (Cherbas et al., 2011) by clustering analysis using Pearson correlation co-efficient scores. A matrix of Pearson correlation scores shown. On the heatmap key (right-hand side), a value of 1 indicates complete correlation; 0 indicates no correlation; −1 indicates complete anti-correlation.

**Supplementary Figure 2:**
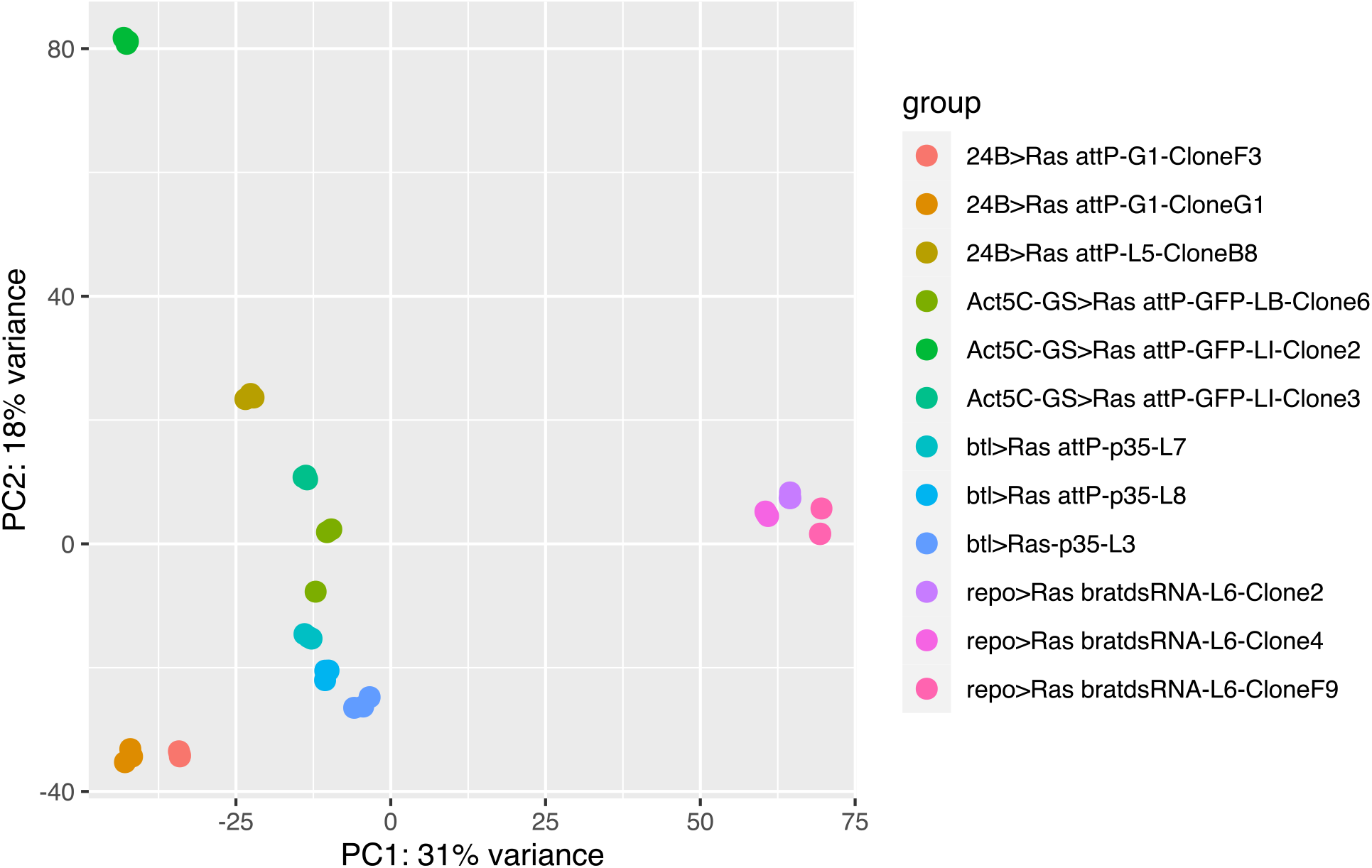
Principal component analysis (PCA) of RNAseq data from the lineage-restricted Ras cell lines. The RNAseq datasets for all cell lines were aligned to the Drosophila genome (r6.48) using STAR. The PCA plot was generated using heatmaply. As expected, cells derived using the same GAL4 driver tend to cluster with one another.

**Supplementary Figure 3:**
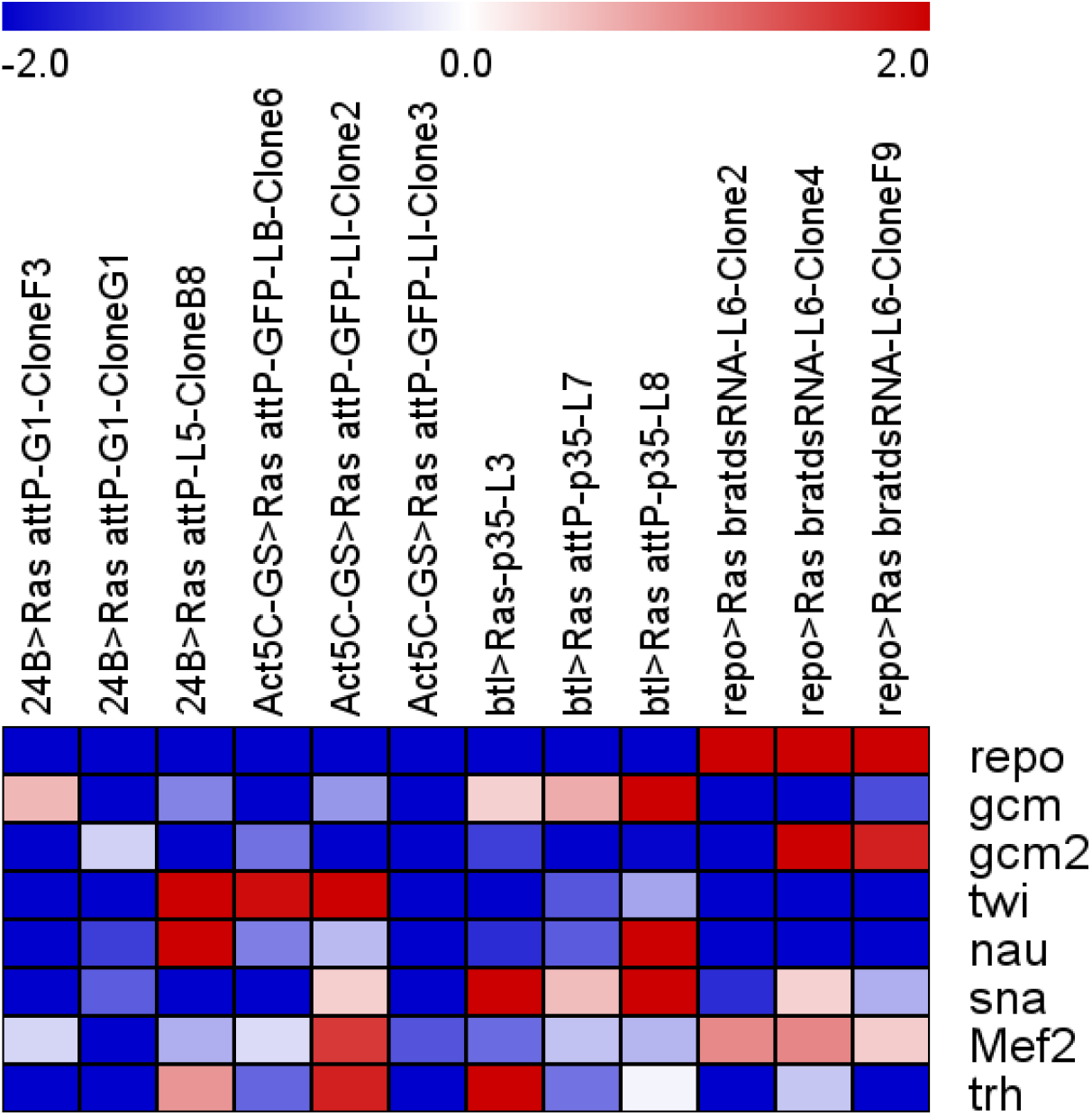
Relative expression of transcription factors associated in the literature with specific tissue lineages in the lineage-restricted Ras cell lines. The set of tissue-specific transcription factors were selected manually from the literature. The fold change in gene expression levels for each cell line as compared to all other cell lines derived using different drivers were calculated using DESeq2 and fold change values are visualized on the heatmap.

**Figure S4.**
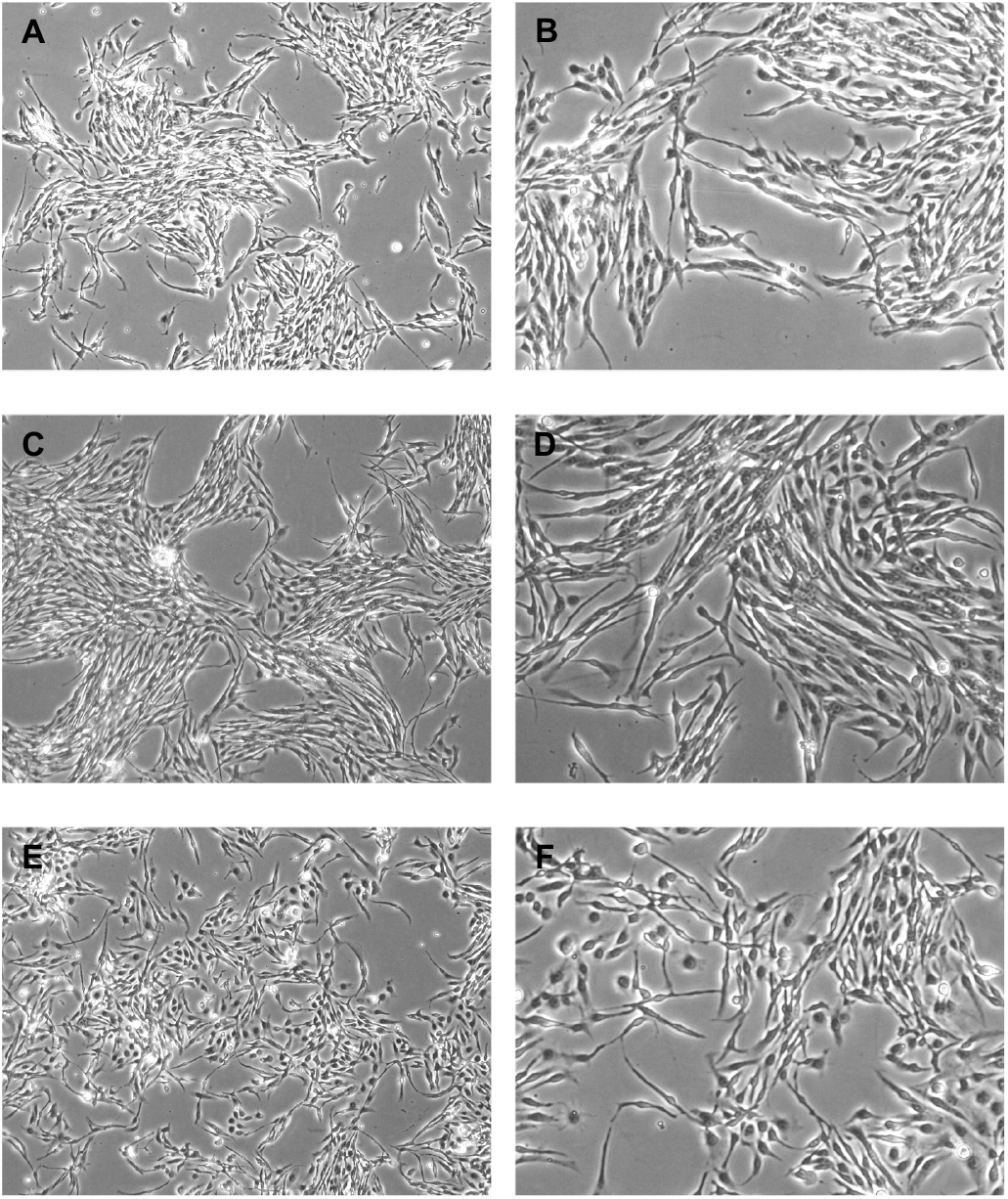
Morphology of Rbr6 clones. (A,B) Rbr6-2. (C,D) Rbr6-2. (E, F) Rbr6-2.

**Fig. S5.**
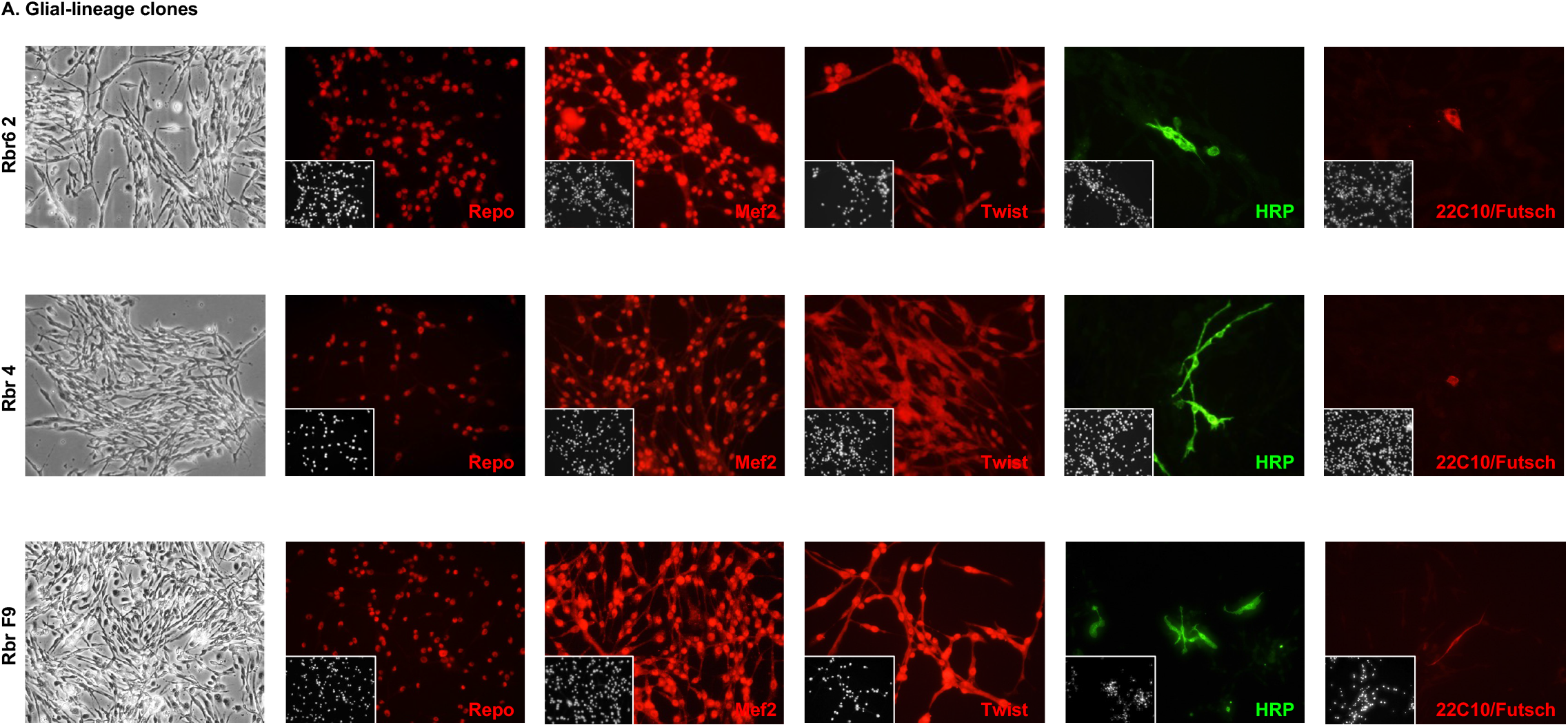
Marker gene expression. **(A) Glial-lineage clones**. Clones were derived from parental line *Repo-Gal4; RasV12; bratdsRNA* Line 6. For each clone, a phase image and marker gene expression are shown. Inset: DAPI, DNA.

**Fig. S5.**
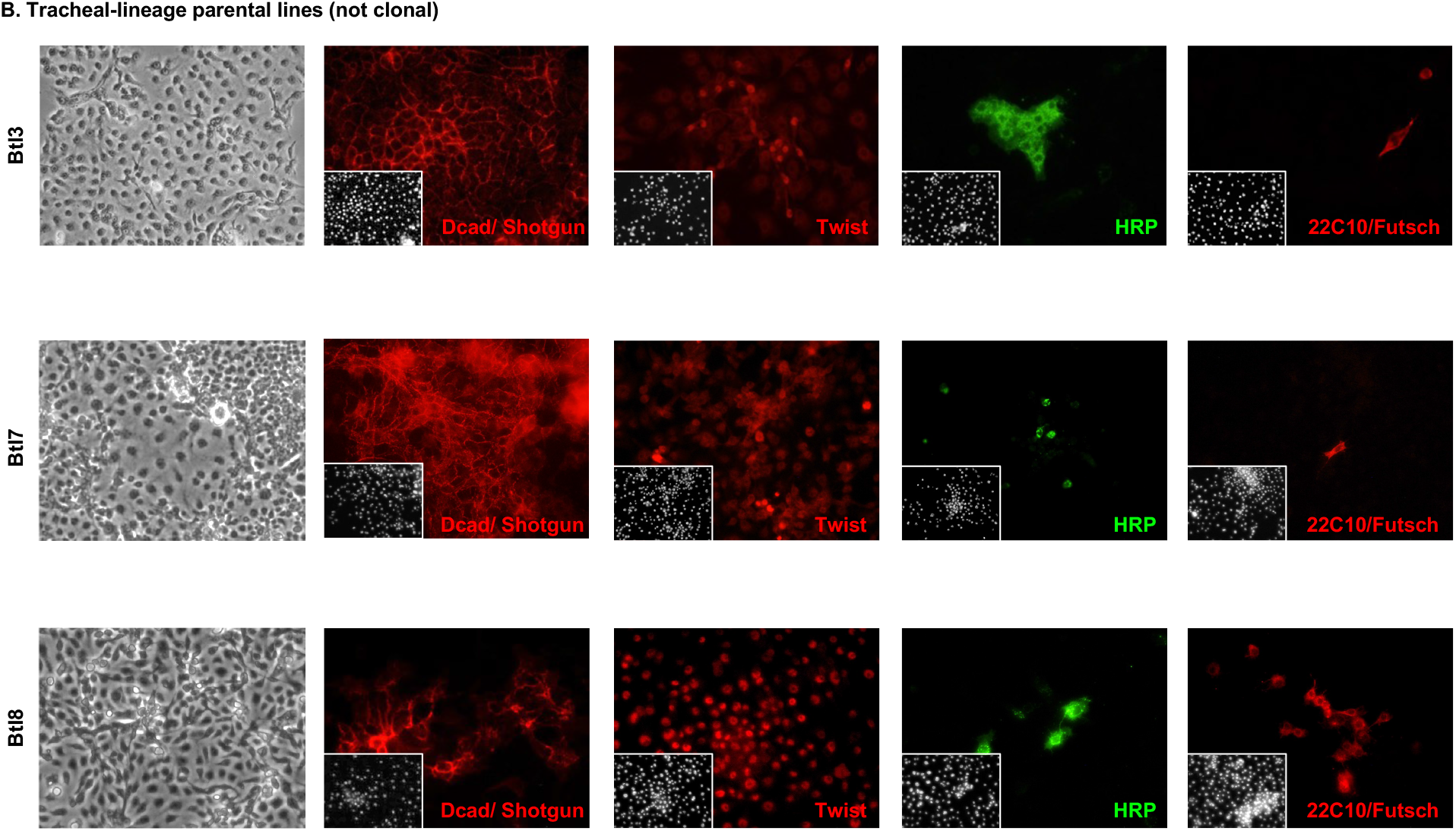
Marker gene expression. **(B) Tracheal-lineage lines**. For each line, a phase image and marker gene expression are shown. Inset: DAPI, DNA.

**Fig. S5.**
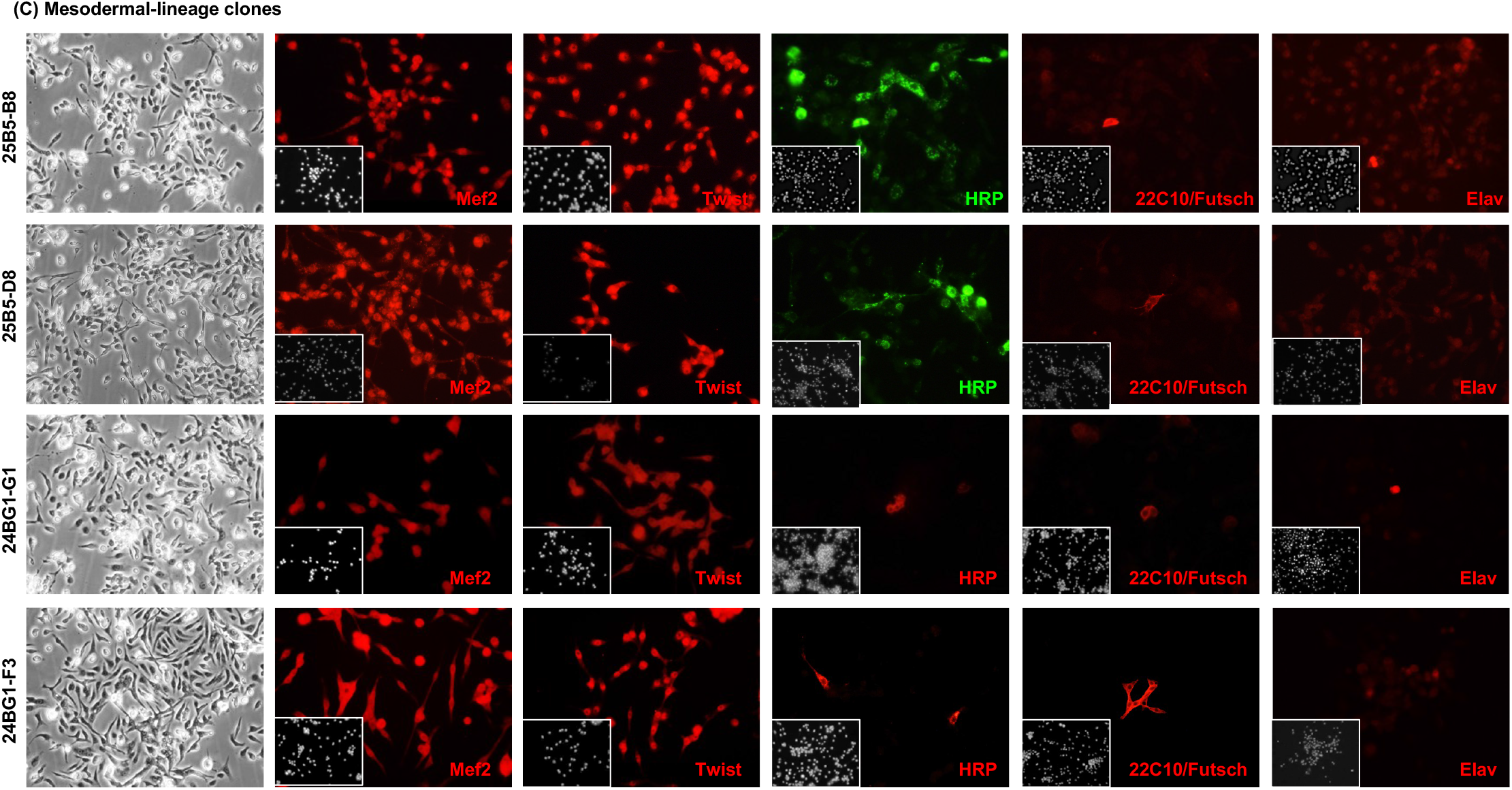
Marker gene expression. **(C) Mesodermal-lineage clones**. For each clone, a phase image and marker gene expression are shown. Inset: DAPI, DNA.

**Fig. S5.**
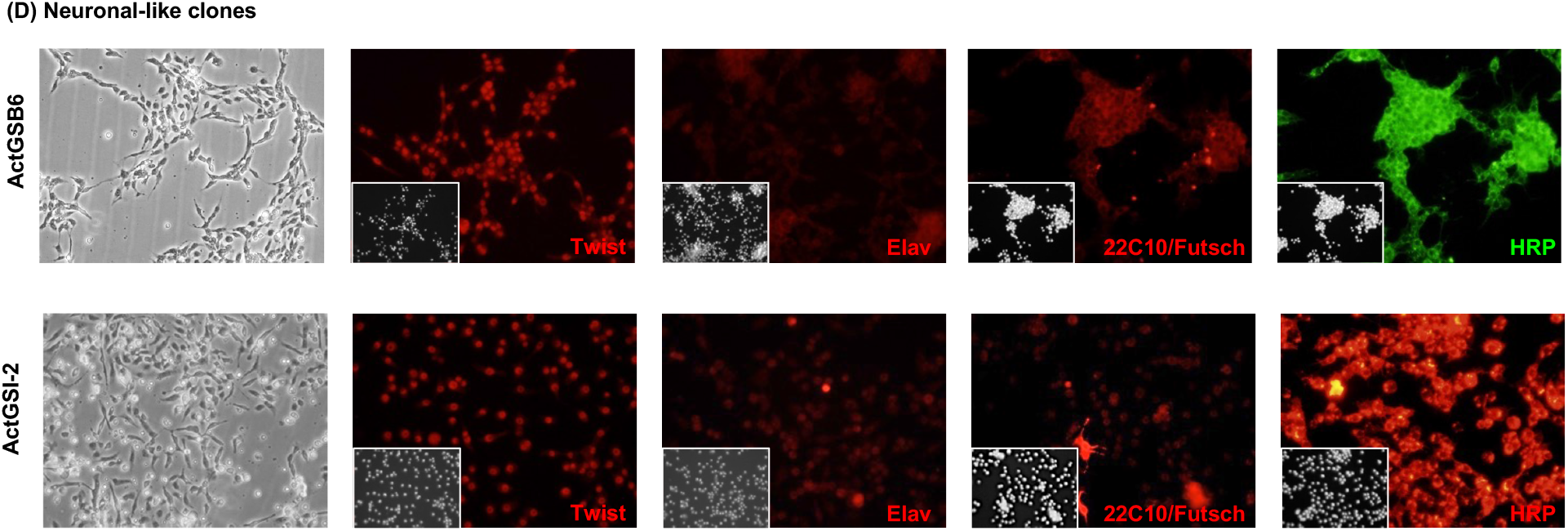
Marker gene expression. **(D) Neuronal-like clones**. For each clone, a phase image and marker gene expression are shown. Inset: DAPI, DNA. Note, the UAS-GFP transgene is not expressed in clone B6.

**Fig. S5.**
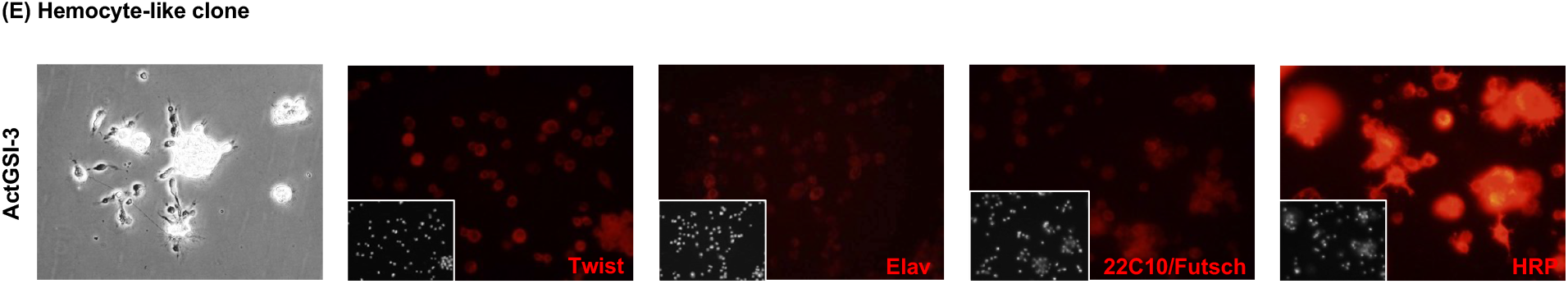
Marker gene expression. **(E) Hemocyte-like clone**. A phase image and marker gene expression are shown. Inset: DAPI, DNA.

**Fig. S6.**
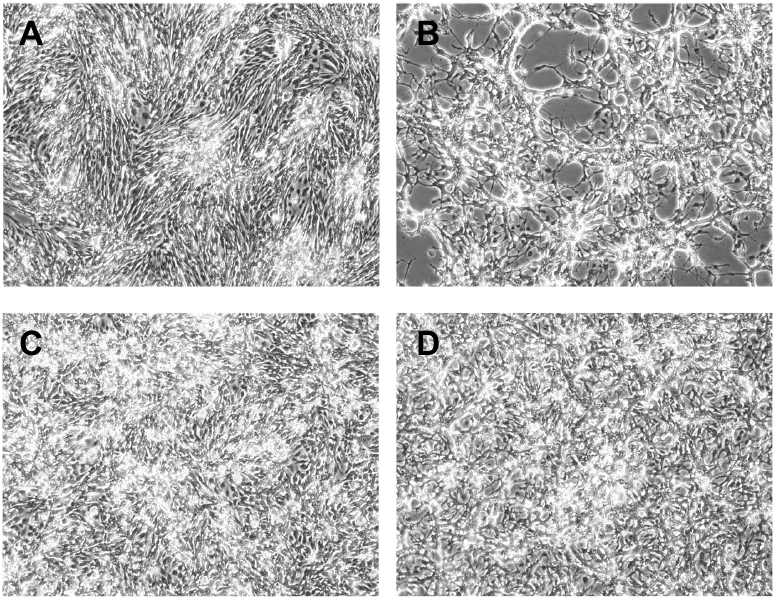
Glial cell morphology with and without ecdysone treatment. Control: A, C. Ecdysone: B, D, cells are shown on day 4, one day after ecdysone-treated cells have received two doses of ecdysone separated by one day. (A,B) Glial clone RBr6-4. After ecdysone treatment, cells appear to fuse and make a lace-like network. (C,D) Glial clone Rbr6-F9. After ecdysone treatment, cells appear similar to control cells.

**Fig. S7.**
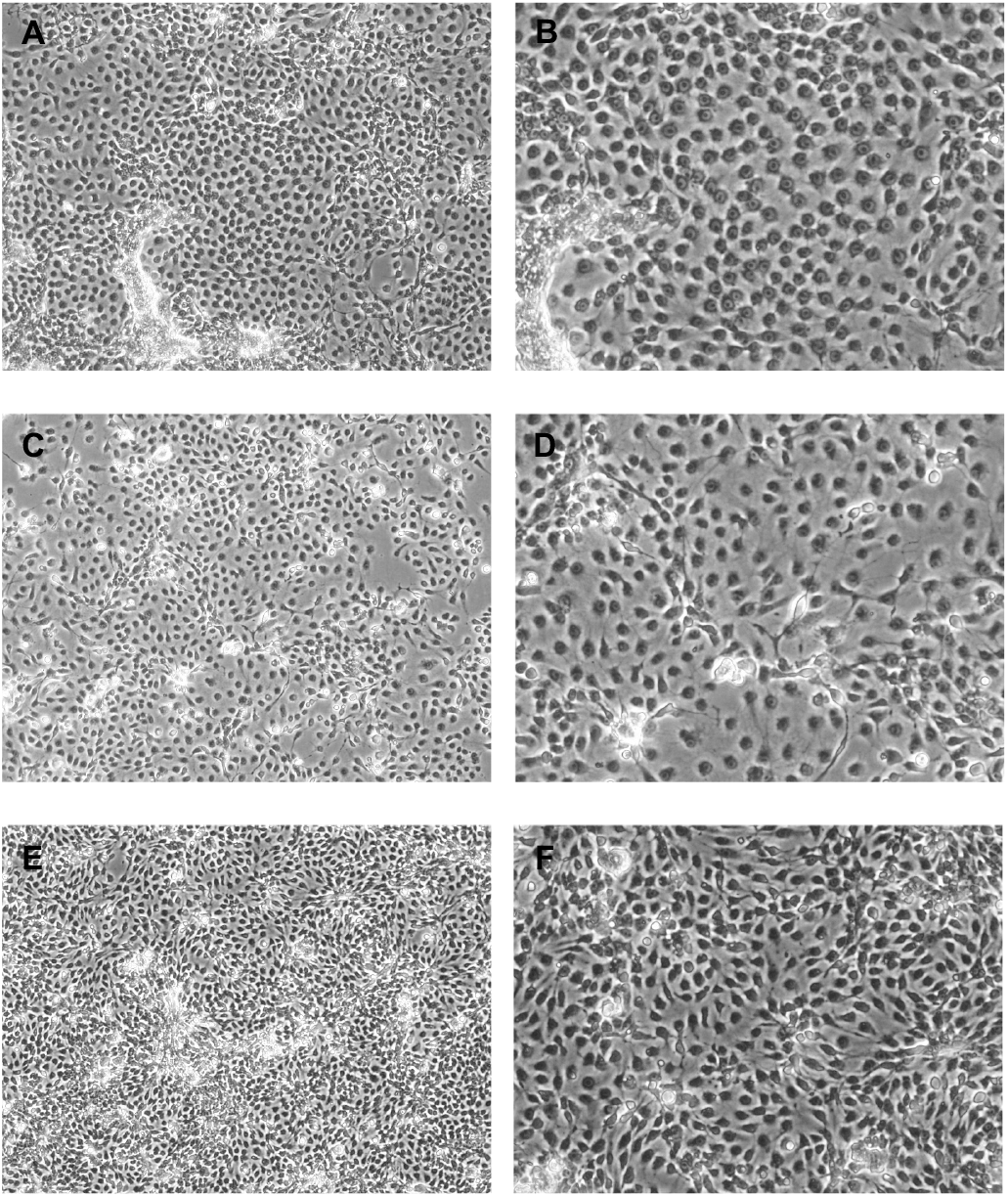
Morphology of tracheal epithelial lineage parental lines. (A,B) Btl3. (C,D) Btl7. (E,F) Btl8.

**Fig. S8.**
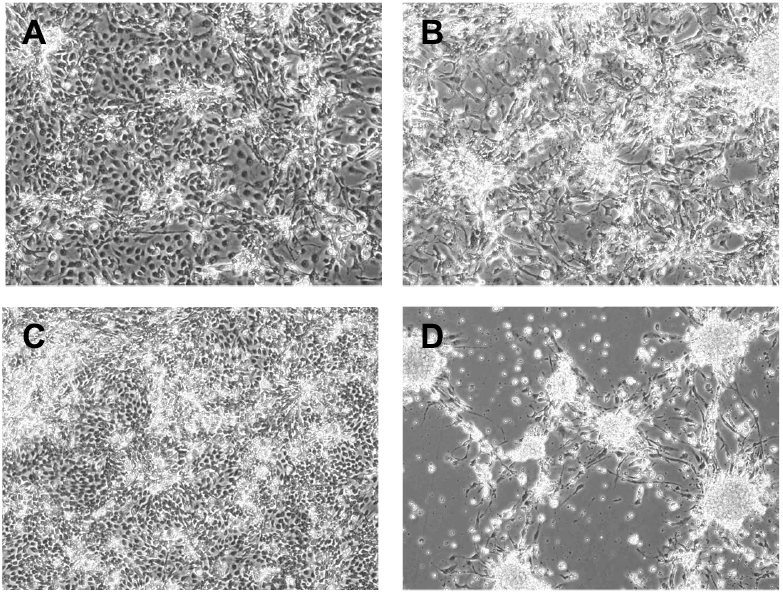
Morphology of tracheal epithelial parental lines after ecdysone treatment. Control: A, C. Ecdysone: B, D. (A, B) Epithelial parental line Btl7. After ecdysone treatment, cells form clusters that extend vertically. (C, D) Epithelial parental line Btl8. There is extensive cell death after ecdysone treatment, suggesting some cells may be of embryonic origin.

**Fig. S9.**
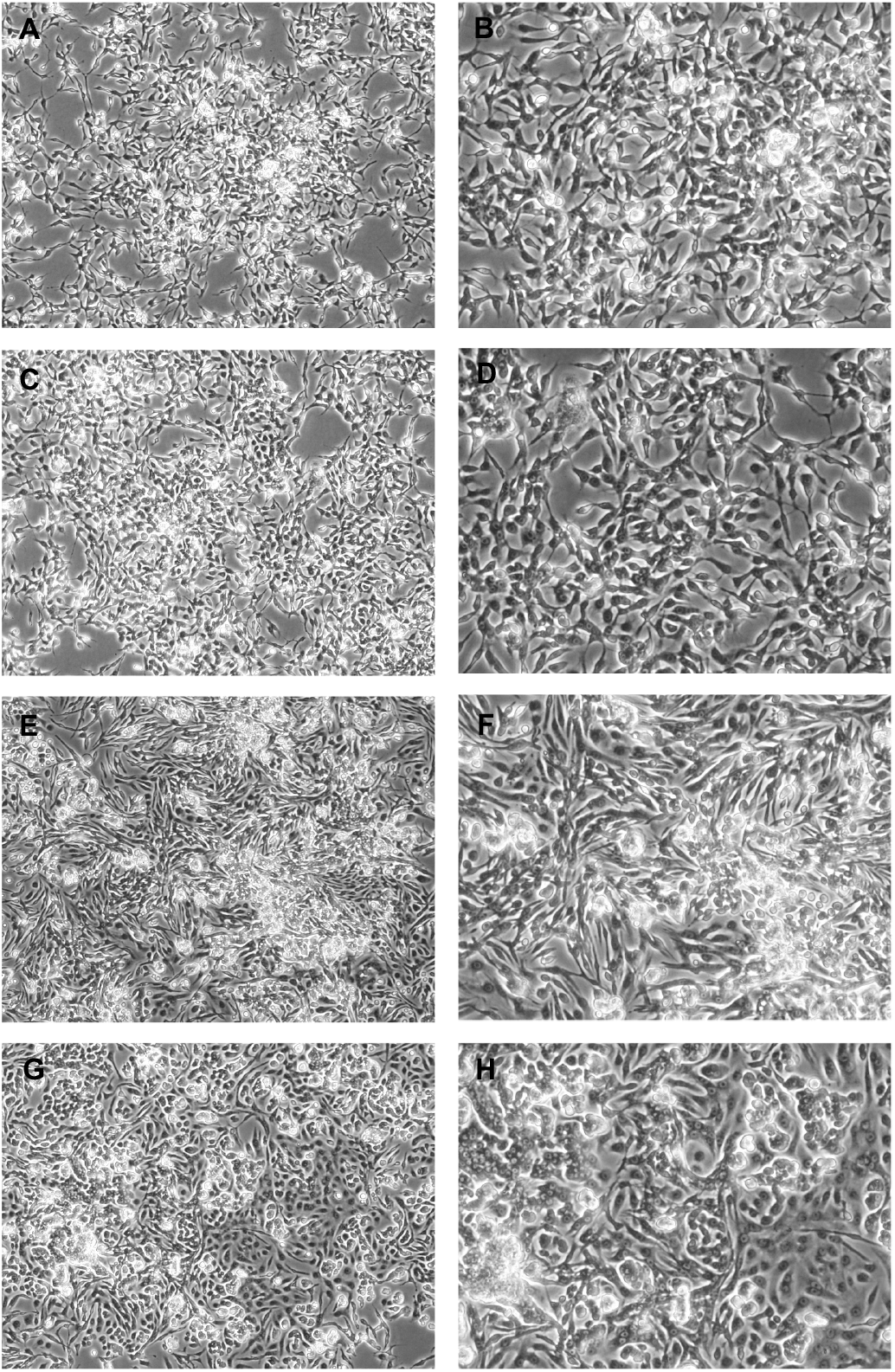
Morphology of mesodermal-lineage clones. (A,B) 24B5-B8. (C,D) 24B5-D8. (E,F) 24BG1-F3. (G,H) 24BG1-G1.

**Figure S10.**
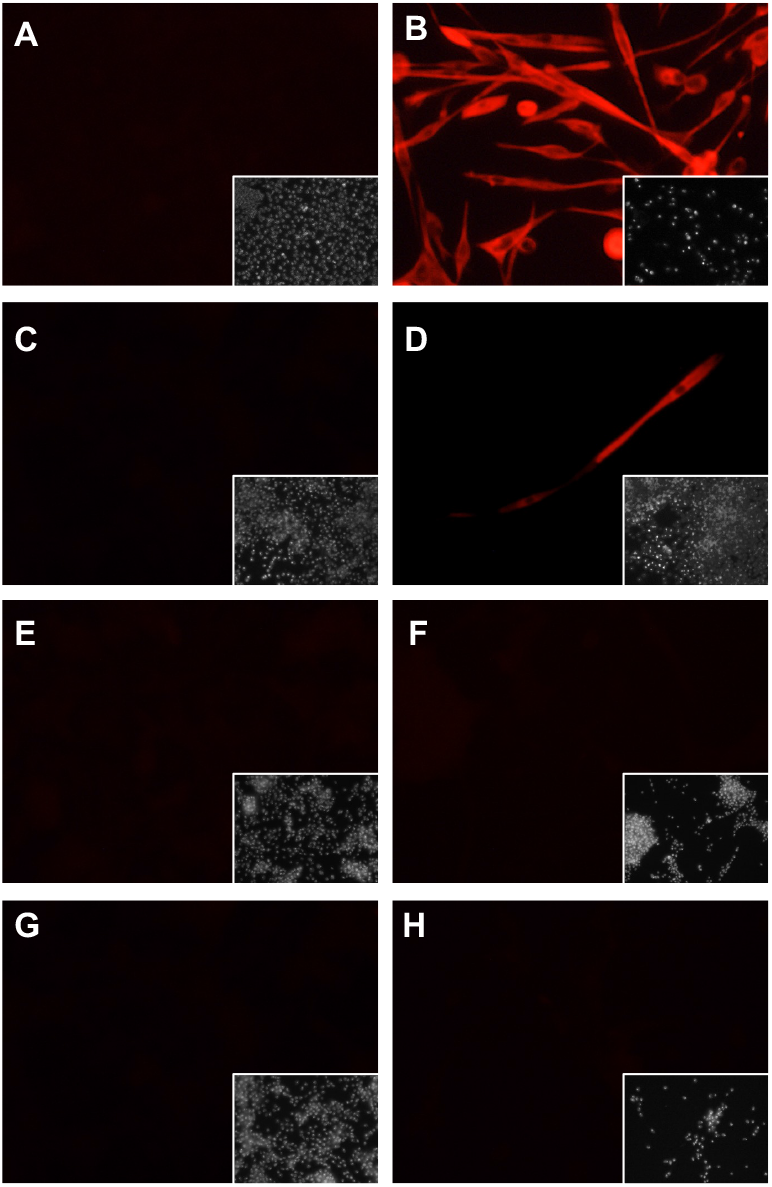
Immunostaining of mesodermal-lineage cells for Myosin heavy chain. (A,C,E,G) Control. (B,D,F,H) Ecdysone. (A,B) Cells of parental line 24BG1 express Mhc after ecdysone treatment at passage 16. (C,D) Rare cells of parental line 24BG1 express Mhc after ecdysone treatment at passage 115. (E,F). Cells of clonal line 24BG1-F3 do not express Mhc after ecdysone treatment. (G,H) Cells of clonal line 24BG1-G1 do not express Mhc after ecdysone treatment. All panels stained for Mhc. Inset: DAPI, DNA.

**Fig. S11.**
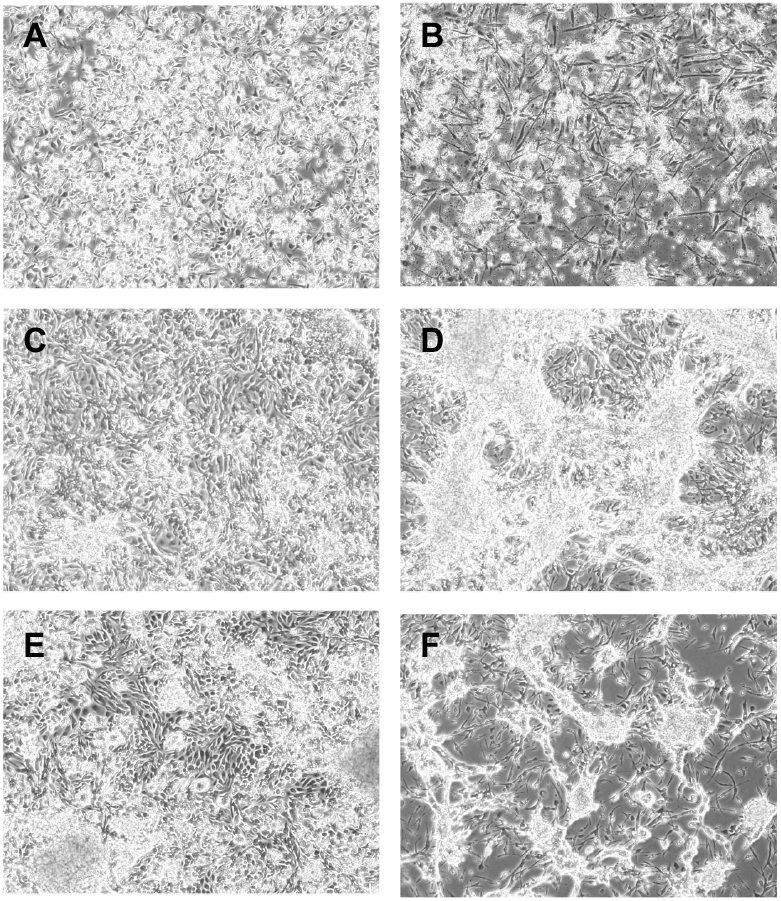
Mesodermal cells showed altered morphology after ecdysone treatment. Control: A, C, E. Ecdysone: B, D, F. (A, B) Mesodermal clonal line 24B5-D8. After ecdysone treatment, cells elongate and begin contraction. (E, F) Mesodermal clonal line 24G1-G1. After ecdysone treatment, cells aggregate, however there is no contraction. (G, H) Mesodermal clonal line 24G1-F3. After ecdysone treatment, cells aggregate, however there is no contraction.

**Fig. S12.**
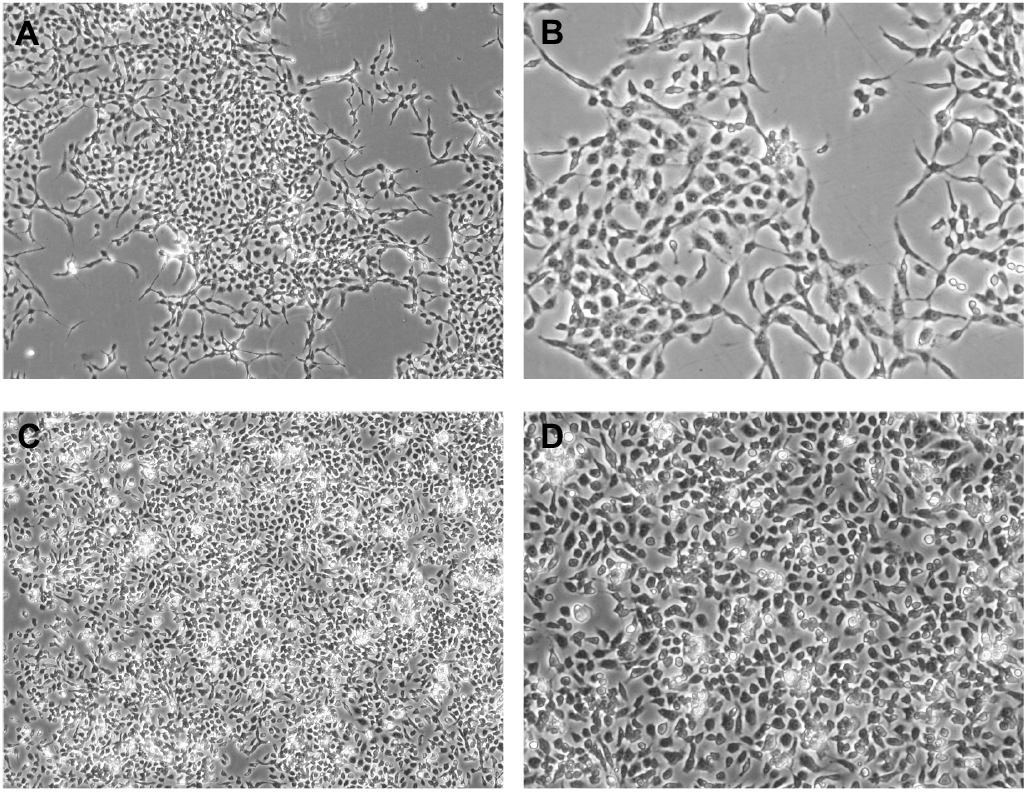
Morphology of neuronal-like clones. (A,B) ActGSB-6. (C,D) ActGSI-2.

**Fig. S13.**
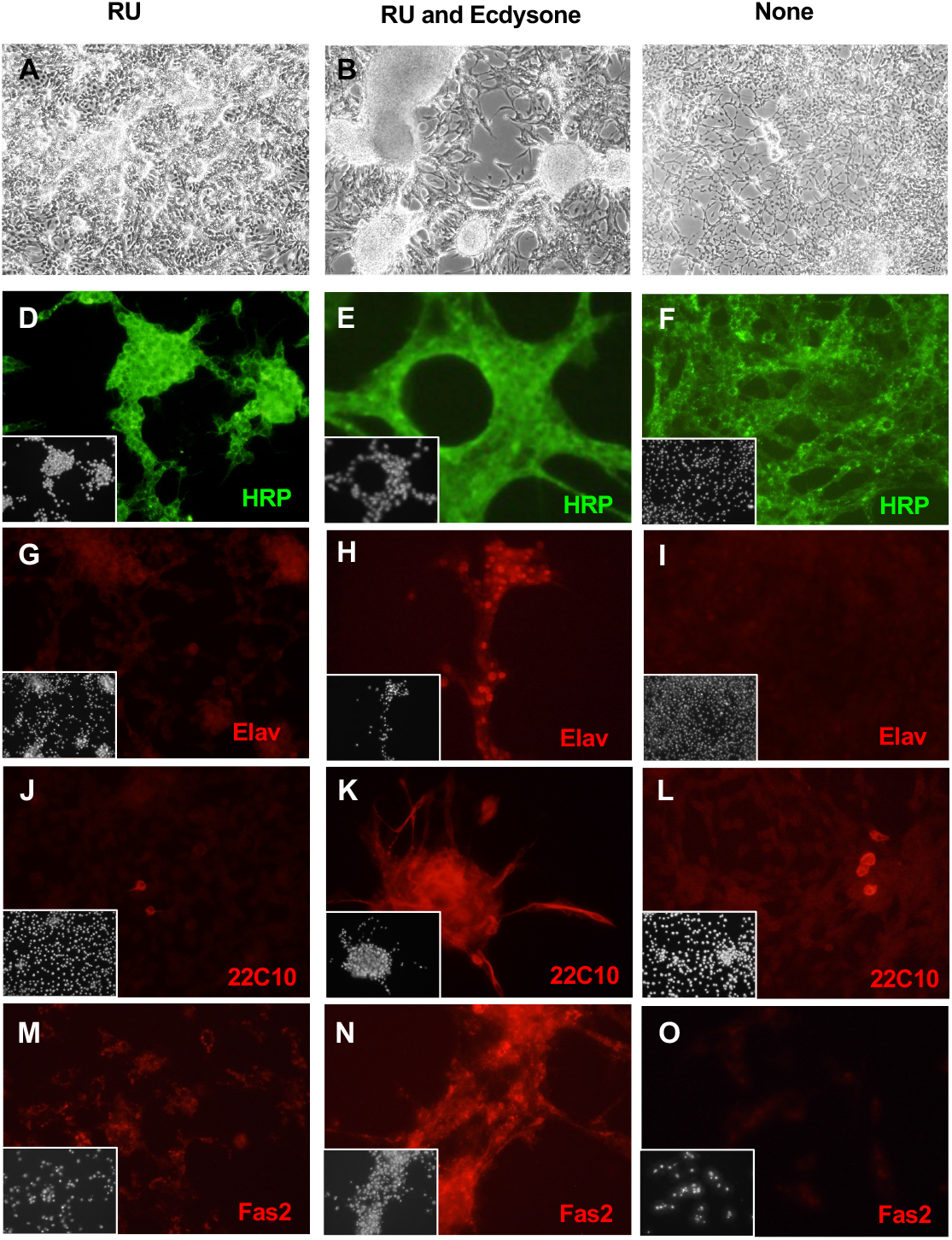
Neuronal-like clone ActGSB-6. Cells were grown in three conditions: RU486 (A,D,G,J,M); RU486 and ecdysone (B,E,H.K,N), or with no additives (C,F,I,L,O). RU486/mifepristone is required for GeneSwtch-Gal4 activation, UAS-Ras_V12_ expression, and cell proliferation. (A) In the growing condition, cells form a sheet, and some cell piles are seen. (B) After ecdysone treatment cells aggregate and form processes. (C) In the quiescent state (no RU), cells cease proliferation and do not reach confluence. (D,E,F) HRP (pan-neural marker) Cells in all conditions are positive for HRP. (G,H,I) Elav (post-mitotic neurons). Elav is elevated after ecdysone treatment (H). (J,K,L) 22C10/Futsch (neuronal morphology). Cells show elevated expression after ecdysone treatment (K). (M.N,O) Fas2 (neural-adhesion protein). Cells show elevated expression after ecdysone treatment (N). Note the UAS-GFP transgene is not expressed in ActGSB-6 cells.

**Fig S14.**
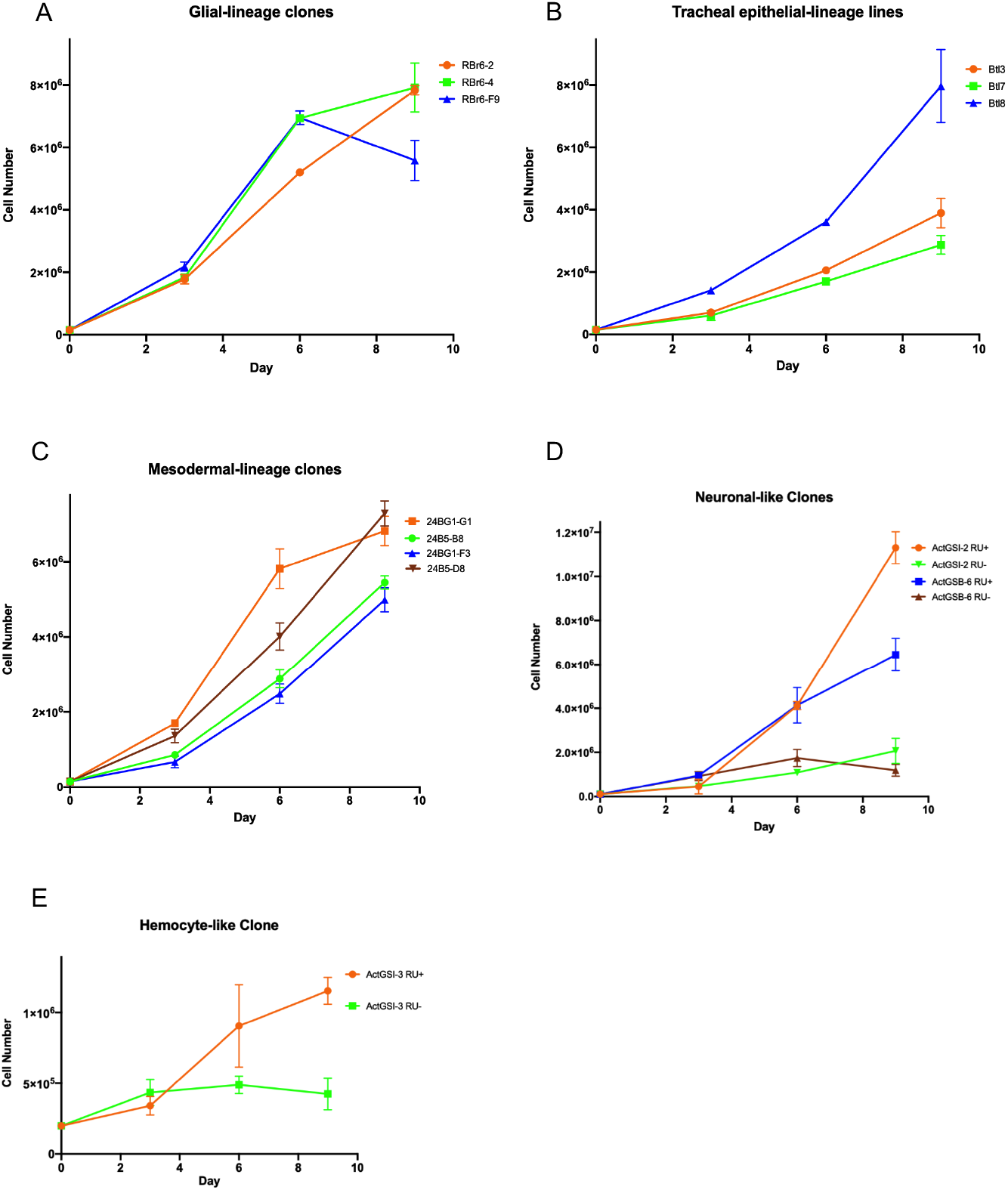
Growth curves. The cells were seeded at 150,000 cells (A,B,C), 100,000 cells (D), or 200,000 cells (E) in 12-well plates and counted on days 3, 6, and 9. (A) Glial lineage clones Rbr6-2, Rbr6-4 and Rbr6-F9. (B) Tracheal epithelium lineage parental lines Btl3, Btl7 and Btl8. (C) Mesodermal lineage clones 24B5-B8, 24B5-D8, 24BG1-F3 and 24BG1-G1. (D) Neuronal-like clones Act5GSI-2 and ActGSB-6. (E) Hemocyte-like clone ActGSI-3. RU-, no drug added (Ras-OFF); RU+, RU486/mifepristone added (Ras-ON).

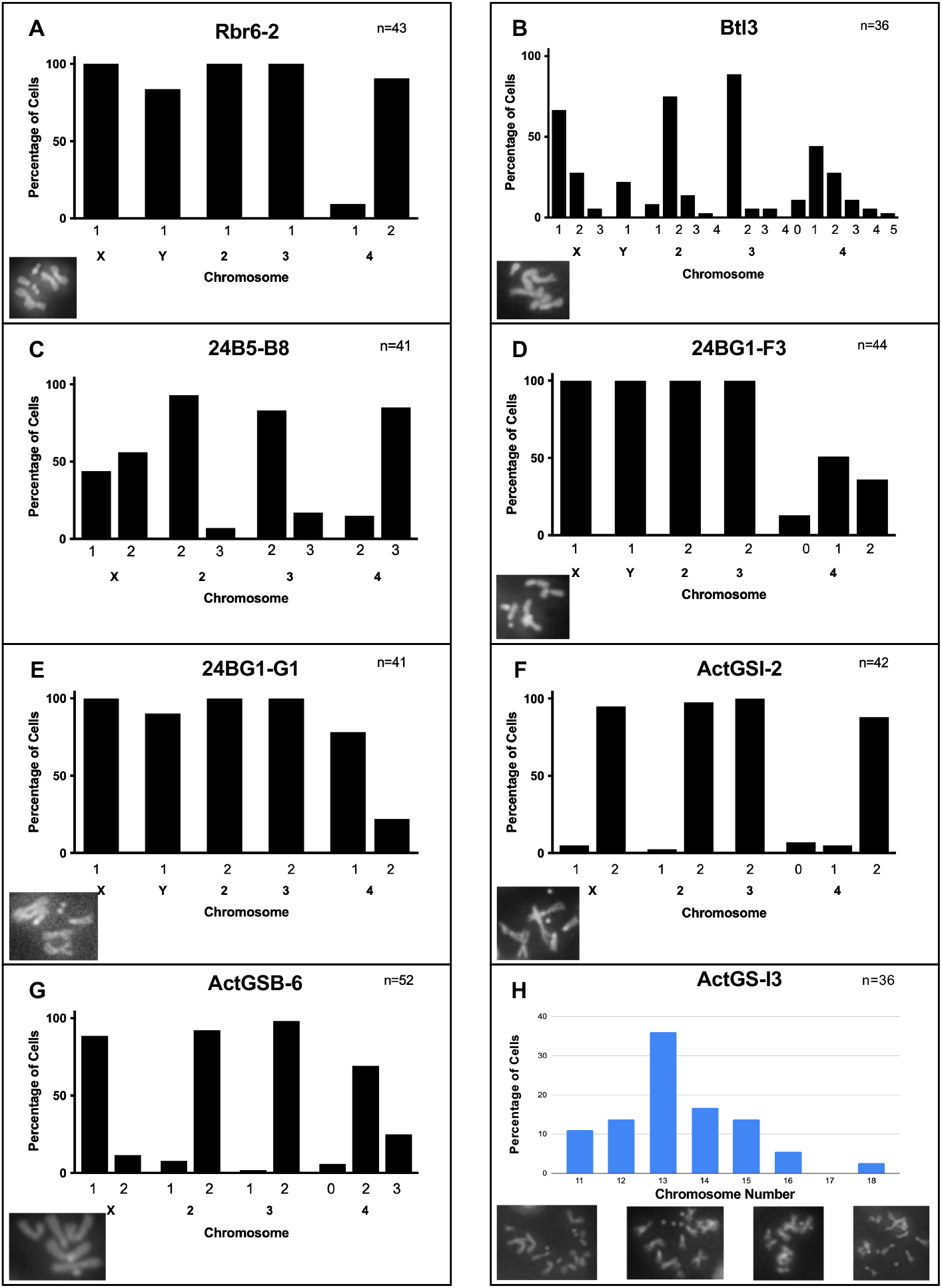

## Notes

### Competing Interest Statement

The authors have declared no competing interest.

